# The N-terminal domain of Eukaryotic Initiation Factor 4B Drives Yeast Translational Control in Response to Urea

**DOI:** 10.1101/819672

**Authors:** Xiaozhuo Liu, Houtan Moshiri, Qian He, Ansuman Sahoo, Sarah Walker

## Abstract

The yeast eukaryotic initiation factor 4B binds the 40S subunit in translation preinitiation complexes (PICs), promoting mRNA binding. Recent evidence suggests mRNAs have variable dependence on eIF4B, suggesting this factor could promote changes in mRNA selection for translation, in order to adapt to stressors. However, the importance of eIF4B and its constituent domains for mRNA selection under diverse cellular and environmental conditions remain undefined. Here we compared the effects of disrupting eIF4B RNA- and ribosome-binding motifs under ~1400 growth conditions. The RNA-Recognition Motif (RRM) was dispensable for stress responses, but the 40S-binding N-terminal Domain (NTD) promoted growth in response to various stressors. In particular, the NTD conferred a strong growth advantage in the presence of urea. Ribosome profiling revealed that the NTD promoted translation of mRNAs with long and highly structured 5-prime untranslated regions, both with and without urea exposure. Our results suggest eIF4B controls mRNA loading and scanning as a part of the PIC, rather than by activating mRNPs prior to ribosome binding. Furthermore, our data indicate the yeast response to urea includes a translational component, driven by production of proteins associated with the cellular periphery. Together our analyses suggest general eIFs can promote diverse cellular responses.

## INTRODUCTION

Translation initiation begins with the formation of a translation preinitiation complex (PIC) comprised of an initiator Met-tRNA·eIF2·GTP ternary complex bound to the 40S ribosomal subunit along with eIFs 1, 1A, 5 and the multisubunit eIF3. Simultaneously, mRNAs are complexed with the eIF4F complex and Ded1/Ddx3, which are proposed to unwind secondary structure and promote PIC binding to mRNA. The eIF4F complex is comprised of eIF4E, which binds the 5’cap structure; eIF4G, a scaffold that binds to mRNAs, eIF4E, eIF4A, and other proteins; and eIF4A, a DEAD-box RNA helicase (1). This complex is thought to serve multiple purposes: 1) interactions of the 5’cap bound to eIF4E with other components of the PIC bound to eIF4G direct PIC loading to the 5’end of mRNAs, and 2) helicase activity of eIF4A melts mRNA secondary structure near the cap and throughout the 5-prime untranslated region (5’UTR) to allow effective loading at the cap and scanning through 5’UTRs. The associated protein eIF4B promotes the activity of the eIF4F complex (2,3).

A number of observations indicate the importance of eIF4B in translating structured mRNAs and promoting the activity of eIF4A/eIF4F both in vitro and in vivo (4–6). In fact, eIF4B in yeast was first discovered by two groups as both a multicopy suppressor of a temperature-sensitive eIF4A mutation, as well as a protein that interacted with antibodies against the 5’-cap complex (7,8). This indicates important functional interaction between eIF4F and eIF4B. One model for eIF4B function suggests that eIF4B enhances eIF4A mRNA helicase activity to allow structured mRNA translation at the step of mRNA activation or scanning (2,9,10). However, recent work indicates that some classes of mRNAs have a hyperdependence on eIF4B while showing less dependence on eIF4A. This suggests eIF4B performs both eIF4A-dependent and eIF4A/eIF4F-independent activities during translation (6,11,12). These eIF4A-independent functions may stem from the ability of eIF4B to bind to the 40S subunit and promote conformational changes in the mRNA binding channel (6,13).

Yeast eIF4B can be divided into four functional domains: an N-terminal domain (NTD), an RNA-recognition motif (RRM), a 7-repeats domain, and a C-terminal domain (Figure 1A; (7,8). While the RRM has a defined globular structure with conserved RNA-binding motifs (14), the other three domains are predicted to be disordered (by analysis of the S288C eIF4B sequence in PONDR (15)). Our previous work demonstrated that eIF4B binds directly to the 40S subunit using both the NTD and 7-repeats domains of the protein, independently of the RRM (13). The 7-repeats domain consists of imperfect repetitions of a ~26 amino acid motif that can also bind directly to single-stranded RNA (7,8,13,16). Binding the 40S by either the NTD or 7-repeats domain promotes the movement of a ribosomal protein, Rps20/uS10 (13). This induces changes in the conformation of multiple rRNA residues on both the solvent and subunit interfaces of the 40S near domains of Rps20/uS10 that reach into the mRNA binding channel. Both the NTD and the 7-repeats domain of eIF4B enhance its affinity to the 40S and are required for robust mRNA recruitment to the PIC, suggesting these ribosome binding activities are critical for translation. Deletion of either domain resulted in decreased rates of mRNA recruitment and translation. However, the defect conferred by deleting the 7 repeats was partially rescued by increasing the concentration of the Δ7-repeats variant in vitro or overexpressing the protein in cells. The heightened concentration of the Δ7-repeats mutant required for maximal rate suggests the repeats must interact with the 40S or another binding partner for maximal affinity and activity. In contrast, increasing concentrations of the Δntd protein did not rescue the decreased rate, suggesting this domain affects the mechanism by which eIF4B accelerates mRNA recruitment (13). Deletion of the NTD, but not other eIF4B domains also confers a dominant-negative overexpression phenotype in cells harboring temperature-sensitive mutant eIF4F alleles, repressing growth even under permissive temperature (17). This suggests the NTD is needed to activate eIF4F. Deletion of either the NTD or 7-repeats domain also increases the amount of eIF4A needed to achieve maximal rate of mRNA recruitment to the PIC, suggesting both domains are needed for optimal functional interaction of eIF4A with the preinitiation complex (13). Evidence was recently provided for RNA-dependent interaction of the 7-repeats domain with eIF4A in vitro (18), and for an RNA-independent interaction of eIF4B with eIF4F in vitro and in vivo (19). These observations together suggest a model in which the 7 repeats domain of eIF4B binds the preinitiation complex, where the NTD can effectively promote eIF4A activity or a conformation of the PIC that allows effective loading and scanning of mRNAs.

**Figure 1.**
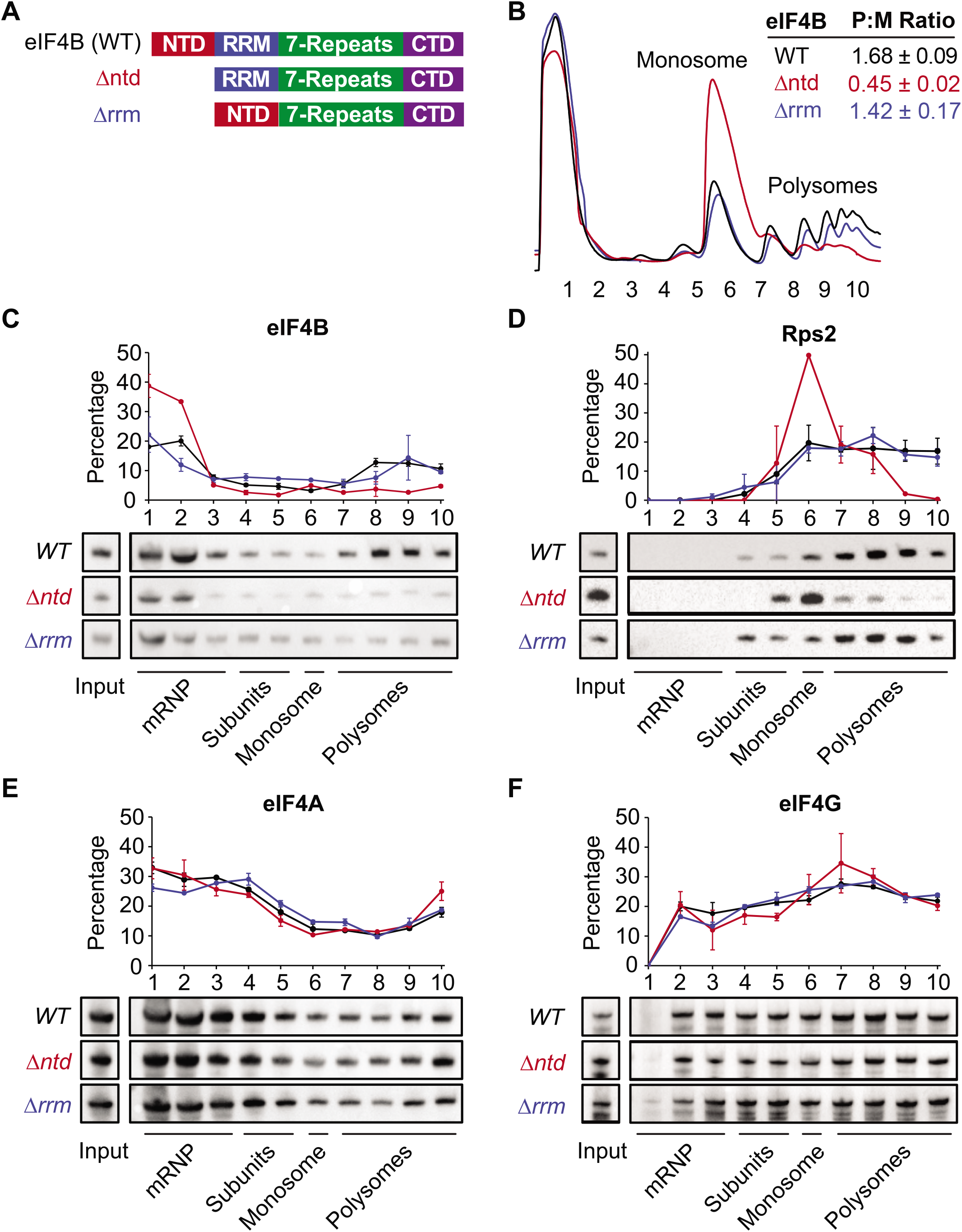
The N-terminal domain of eIF4B binds the ribosome and promotes translation in a prototrophic yeast strain. **A.** Schematic of eIF4B functional domains in WT, Δntd, and Δrrm constructs. **B**. Lysates from strains harboring WT (black), Δntd (red), or Δrrm (blue) eIF4B were fractionated on 10-45% sucrose gradients and monitored by absorbance at 254 nm. Polysome to monosome (P:M) ratios were calculated by quantifying the area under the curve for the indicated monosome and polysome peaks. The Mean P:M ratios from 3 or more biological replicates (representative traces shown) are reported +/- standard error of the mean. **C-F**. Lysates were crosslinked with 1% formaldehyde prior to running on 10-45% sucrose gradients. The distributions of eIF4B (C), Rps2 (D, an indicator of 40S subunits/monosomes/polysomes), eIF4A (E), and eIF4G (F) in mRNP, subunit, monosome, and polysome fractions were determined by western blotting indicated gradient fractions. The line graphs show the mean percentages of total indicated protein (quantified across the gradient) that was present in each fraction.

In contrast to the 40S binding activities of the NTD and the 7 repeats domains, disrupting the RNA-binding activity of the RRM of yeast eIF4B did not affect growth or translation activity in vitro or in vivo, unless combined with other mutations that affect function on their own, such as deletion of the NTD or 7-repeats (13). This suggests that the RNA-binding RRM can stimulate the required function of the NTD and 7 repeats, although to a limited extent which does not accelerate growth rate under standard laboratory conditions. The RRM is the only large functional domain with considerable conservation between yeast and human eIF4B (20), although human eIF4B harbors small motifs of homology to the NTD and a segment of homology to the core sequence of a single yeast repeat just downstream of the RRM. These motifs in the human factor are sufficient to rescue function of yeast eIF4B variants lacking the analogous motifs (17).

Here we investigated the role of the RNA-binding RRM and the 40S-binding NTD of eIF4B on translation and growth in response to stress conditions. We found that the NTD allowed enhanced growth in conditions that require robust cellular integrity, including media containing 3% w/v urea. Deletion of the NTD resulted in reduced eIF4B association with ribosome fractions, and large decreases in translation that were exacerbated by addition of urea. Analysis of the structural content of mRNAs that depended on the NTD support the model that eIF4B is necessary to enhance translation of mRNAs with long, structured 5’UTRs that showed less enrichment with closed-loop factors (6). eIF4B interaction with translating PICs was disrupted by truncation of the NTD, while interaction with 40S fractions was actually enhanced, suggesting this domain enhances steps downstream of 43S formation, such as PIC scanning. The mRNAs that responded to eIF4B NTD-deletion encode cell wall and membrane proteins, as well as proteins involved in ER/Golgi. Further analysis of gene ontology classes for mRNAs that were highly dependent on the eIF4B NTD showed that mRNAs encoding membrane and trafficking proteins, irrespective of eIF4B NTD-dependence had more structured 5’UTRs than other yeast mRNAs. This is expected to allow a high capacity for translational control of membrane proteins in response to various stressors that challenge cellular integrity. Together our data suggest eIF4B drives changes in translation of mRNAs encoding membrane and cell wall proteins to allow cells to adapt to diverse cellular environments.

## MATERIAL AND METHODS

### Construction of yeast strain and plasmids

Yeast strains (Table S1) YSW3 and YSW4 were generated by tetrad dissection of strain FJZ001 (*MATa/MATα, his3Δ1/his3Δ1 leu2Δ0/leu2Δ0 met15Δ0/MET15 LYS2/lys2Δ0 ura3Δ0/ura3Δ0 TIF3/tif3Δ::hisG-URA3-hisG*) (13), followed by isolation of LYS+, HIS-, LEU-, and MET-haploid clones, and finally counterselection on 5-FOA for removal of the *URA3-HisG* cassette of the URA+ *tif3*Δ strain. *TIF3* alleles with the native promoter and terminator encoding C-terminal hexahistidine tagged-eIF4B or variants were Gibson-cloned (NEB, USA) into the BamHI site of single-copy vector pHLUM (21). DNA sequences of the entire PCR-amplified regions were verified in the assembled plasmids (Table S2). Plasmids were transformed into YSW4 to generate prototrophic strains. Transformed strains were verified by Western blot for correct eIF4B variant expression (Figure 2D, Table S1).

**Figure 2.**
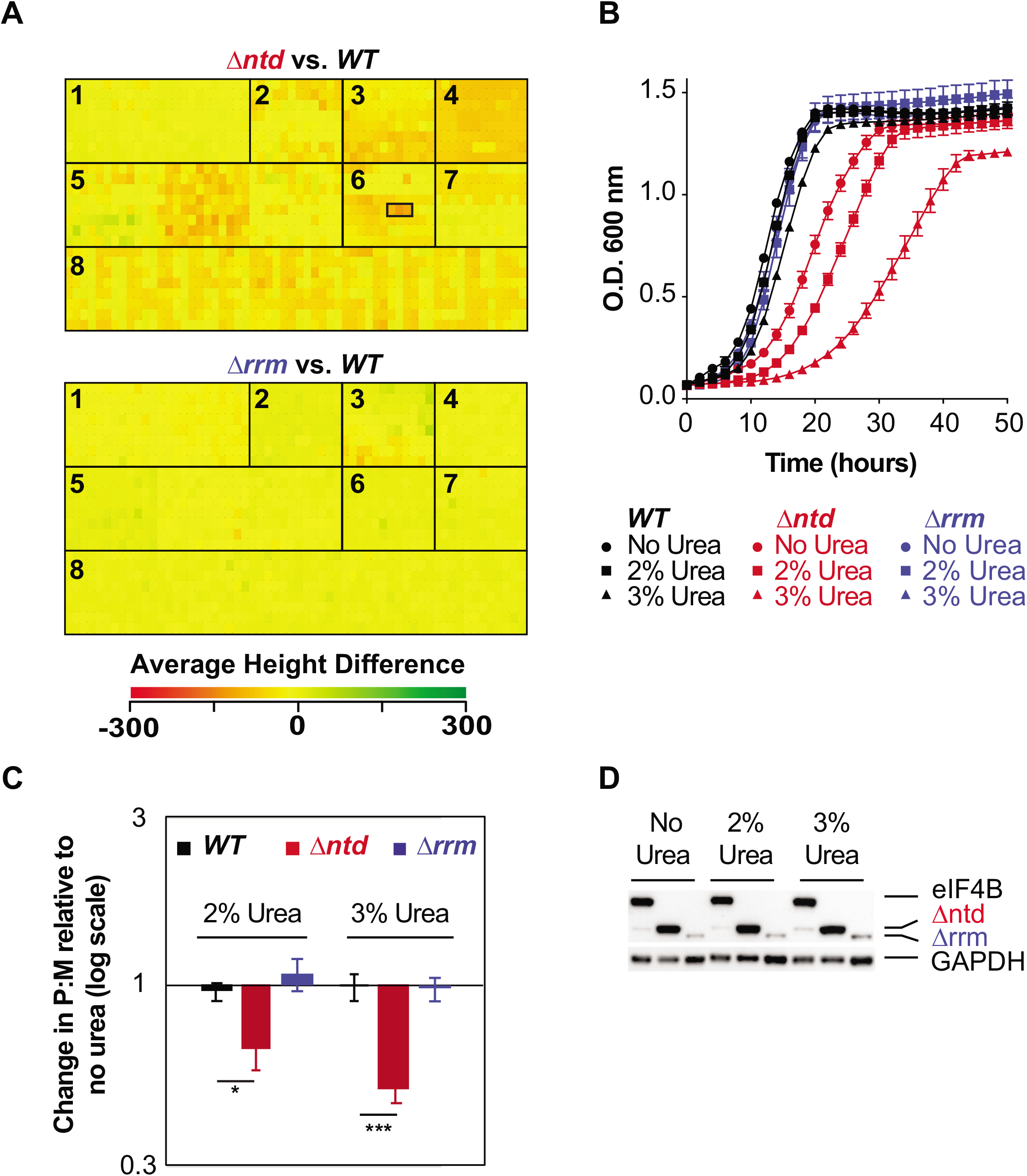
eIF4B binding to the 40S promotes resistance to stressors that challenge cellular integrity through changes in translation. **A.** Heatmap of average height differences for cellular fitness between the NTD (top) or RRM (bottom) deletion mutant (green) and WT (red) under 1440 metabolic and chemical conditions as assayed by Phenotype microarray, which includes seven panels in 15 96-well plates: 1. Carbon Sources; 2. Nitrogen Sources; 3. Phosphorus and Sulfur Sources; 4. Nutrient Supplements; 5. Peptide Nitrogen Sources; 6. Osmolytes; 7. pH; and 8. Chemical Sensitivity. Wells containing 2%, 3%, and 4% urea are outlined by a black box, left to right, for Δntd. 3% Urea gave the largest difference in WT and Δ*ntd* fitness. There were no conditions that gave a significant change for Δrrm **B.** Growth curves of WT (black), Δntd (red), and Δrrm (blue) grown with 0% (circles), 2% (squares), or 3% urea (triangles). Results from biological triplicates ± SEM are shown. **C**. Relative change in polysome to monosome (P:M) ratio upon exposure to urea was determined as in 1B with representative traces shown in Fig. S2 in the presence and absence of 3% urea. Biological triplicates ± SEM are shown with P values from Student’s t test indicated (*P < 0.05). **D**. eIF4B levels in WT, Δntd, and Δrrm strains after growth with 0, 2%, or 3% urea in Synthetic Dextrose media.

Plasmids (Table S2) for monitoring translation reporter expression were constructed using the Yeast Mo-Clo Golden Gate cloning system as described (22). The promoter, 5’UTR, and first 30 nucleotides were amplified from yeast (BY4741) genomic DNA and cloned into parts vector pYTK001 via BsaI assembly. An intermediate vector (pAS45) contained the Parts 2-4 (pYTK047) insert for screening *E.coli* colonies by replacement of GFP was combined with the FIG2 and VBA2 gene insertion parts vectors, the C-terminal Venus fusion tag (pYTK045), and a terminator part vector (pYTK053) for BsmBI assembly of the complete reporter vectors. Plasmids were verified by sequencing.

### Phenotyping Experiments

Previous work showed that inclusion on the same plasmid of the four requisite metabolic genes lacking in the parental genome of the barcoded yeast knockout collection (*HIS3, LEU2, URA3*, and *MET17*) provided a selective growth advantage, regardless of whether the metabolites these genes produce were included in the media. Hence, the plasmid with four metabolic markers is effectively retained even in rich growth media without selection, and synthetic growth effects between metabolic gene deletion and the mutation of interest can be avoided (21). Prototrophic strains YSW5, 6, and 7 were sent to Biolog for Phenotype Microarray screening at 30°C in duplicate, using plates 1-10 and 20-25 (Biolog Inc, USA) (23). Plates were read every 30 minutes for 48 hours.

### Yeast Growth

Yeast were cultivated in liquid or on solid (2% agar) Synthetic Drop-out (SD) media (20 g/L glucose, 1.71 g/L yeast nitrogen base without amino acids, 5 g/L ammonium sulfate; Sunrise Scientific Products, USA) at 30°C. For assays with additives, yeast cells from an overnight culture in SD media were diluted to an OD_600_ of 0.05 in SD media or SD media supplemented with additive (e.g. 2% or 3% urea, 1.5 mg/ml caffeine), and allowed to grow to mid-log phase with vigorous shaking at 30°C, unless otherwise noted. Automated growth curves were performed in 96-well plates (200 μl SD per well) by taking OD_600_ measurements every two hours for 48 hours while incubating at 30°C with double-orbital shaking in a Spark plate reader (Tecan, Switzerland).

### Analysis of Polysome:Monosome ratios

Polysome analysis was performed as described previously (13,24). For figure 1B, yeast cells were cultured in SD medium with or without additives as noted at 30°C to an OD_600_ of 1.5. Cycloheximide (Gold biotechnology, USA) was added to a final concentration of 50 μg/ml and incubated for 5 minutes at 30°C with shaking before collecting cells by pelleting in centrifuge bottles packed with ice. Pellets were resuspended in 1/3 of the pellet weight of breaking buffer (20 mM Tris-HCl at pH 7.5, 50 mM KCl, 10 mM MgCl2, 1 mM DTT, 200 μg/mL heparin, 50 μg/mL cycloheximide, and 1 Complete EDTA-free Protease Inhibitor Tablet [Roche]/50 mL buffer), dropped into liquid nitrogen, and lysed in a Nitrogen Mill for 10 Cycles following a precool of 15 minutes with the following parameters: 1 minute run, 2 minutes cool, rate=15 (Spex Sample Prep, USA). 25 A_260_ units of lysates were resuspended in 1.5 volumes of the pellet weight of ice-cold breaking buffer and clarified by spinning at 14,000 rpm for 15 minutes at 4°C prior to separation by velocity sedimentation on 5-45% sucrose gradients (20 mM Tris-HCl [pH 7.5], 50 mM KCl,10 mM MgCl_2_,1 mM DTT, 5-45% sucrose) by centrifugation at 39,000 rpm for 2.5 h at 4°C in a Beckman SW41 rotor. Gradient fractions were separated on a gradient station (BioComp, Canada) while scanning at 254 nm. The areas under the monosome and polysome peaks, determined in GraphPad Prism software, were used to calculate the P/M ratio.

### Analysis of initiation factor association with subunits and polysomes by crosslinking and gradient ultracentrifugation

To monitor association of eIF4B and other proteins with 40S subunits, formaldehyde crosslinking analysis was performed as described (25) with the following differences: Yeast cells were grown to an OD_600_ of ~0.8 in SD media at 30°C. Cultures were poured into bottles packed with ice containing formaldehyde for a final concentration of 2% and incubated for 60 min prior to collection and lysis. Cells were lysed in a nitrogen mill as described above and resuspended to 20 A_260_ units per 300 μl. 300 μl of crosslinked lysates were then loaded and separated by spinning at 41000 rpms for 5 hours on 7.5%-30% sucrose gradients in an SW-41 rotor, and 0.63 ml fractions were collected upon fractionation on a Biocomp gradient station. Fractions 1-2 were combined prior to loading. Fractions up to the 40S peak were analyzed by western analysis. Three biological replicates were performed.

For observing polysome association, Cycloheximide was added to the culture to a final concentration of 50 μg/ml and incubated for 5 minutes at 30°C with shaking before harvesting cells on ice. Cells were lysed and resuspended in BBK buffer as above, then formaldehyde was added to the resuspended lysates at a final concentration of 1%. Crosslinking was carried out for 30 min on ice before quenching with glycine at 0.1M. Crosslinked cell lysates were layered on 5%-45% gradients and 10 1 ml fractions were collected for analysis by western blotting (25). Two biological replicates were performed.

### Western analysis

To verify that eIF4B variants were expressed under stress, yeast cells were grown in SD media with or without 2% or 3% urea, and harvested at an OD_600_ of 1.0. Whole cell extracts (WCEs) were prepared by extraction with trichloroacetic acid (TCA) and subjected to Western blot analysis as described previously (13) using antibody against the His6 epitope (EMD Millipore/Novagen 70796, 1:2000 dilution) Experiments were repeated three or more times from Biological replicates.

For analysis of eIF4B position within gradients, 0.5 ml of each gradient fraction was precipitated by addition of 1 ml 100% Ethanol and spinning for 13 minutes at 13,000 x g, resuspended in SDS loading buffer, and resolved by SDS-PAGE, followed by Western blotting using anti-His antibody. 40S subunit (and 80S/polysome) containing fractions were verified from the same samples by blotting yeast ribosomal small subunit protein Rps2 (Aviva ARP63572_P050, 1:2000 dilution.) Rabbit antibodies to purified recombinant yeast eIF4A (1:20,000 dilution) and eIF4G1 (1:1000 dilution), generated by Invitrogen/Pierce custom antibody services and verified against the recombinant proteins, were used to determine the position of those proteins within gradients (26). Each experiment was repeated three or more times from biological replicates.

To determine changes in FIG2 and VBA2 translation reporter fusions, TCA precipitated lysates were prepared as above from cells grown in media with or without 3% urea or 1.5 mg/ml caffeine. Lysates were separated on a stain-free SDS-PAGE gel (Bio-rad), then blotted with mouse anti-GFP antibody (Roche 11814460001) and anti-mouse-HRP secondary (Cell Signaling 7076). GFP was normalized to total protein bands per lane (visualized by a stain-free scan on a Bio-rad gel doc). Each experiment was repeated at least two times from biological replicates.

### RiboSeq and RNASeq library preparation

Ribosome footprint profiling was conducted as described (27–29) with minor modifications. Yeast cells at an OD_600_ of approximately 0.8 were rapidly harvested by vacuum filtration through a 0.45 μm Whatman cellulose nitrate membrane filter (GE Healthcare Life Sciences, USA) at room temperature by scraping the slurry into liquid nitrogen. Cells were lysed as above in a Nitrogen mill, and thawed and suspended in lysis buffer (20 mM Tris [pH 8], 140 mM KCl, 1.5 mM MgCl_2_, 1% Triton X-100, 100 μg/ml cycloheximide). 25 A_260_ units of extract were treated with 87.5 Units of RNase If (M0243, NEB, USA) for 1h at 22°C on a rotator, then separated on 10-50% sucrose gradients (20 mM Tris [pH 8], 140 mM KCl, 1.5 mM MgCl_2_, 1mM DTT, 100 μg/ml cycloheximide, 10-50% sucrose) by centrifugation at 40,000 rpm for 3 hours at 4°C in a Beckman SW41 rotor and fractionated as above. Ribosome-protected mRNA footprints were purified from the nuclease-treated monosome fraction by addition of SDS to 0.8% at 65°C, followed by extraction with acid phenol [pH 4.5] (Ambion, USA) and then chloroform/isoamyl alcohol extraction. 300 mM NaOAc [pH 5.2] was added to the aqueous phase and RNA was precipitated with one volume of isopropanol before resuspending in 10 mM Tris-HCl [pH 8]. RNA footprints from 25 to 35 nt were size-selected on a 15% denaturing PAGE gel, eluted by crushing and soaking gel fragments in RNA elution buffer (300 mM NaOAc [pH 5.5], 1 mM EDTA, 0.1 U/μl SUPERaseIn (Life Technologies, USA)), and dephosphorylated using T4 Polynucleotide Kinase (M0201, NEB, USA) prior to isopropanol precipitation and resuspension in 10 mM Tris [pH8]). A preadenylated universal linker (5’-rAppCTGTAGGCACCATCAAT-NH2-3’) was prepared in house or purchased from NEB (S1315S) and ligated to the 3’ ends of the dephosphorylated footprints using T4 RNA Ligase 2, truncated (MO242L, NEB). rRNA was depleted using the Yeast Ribo-Zero Gold rRNA removal kit (Illumina, USA). First strand synthesis was performed with Superscript III (Life Technologies, USA) and reverse transcription primer NINI9 (5’-/5Phos/AGA TCG GAA GAG CGT CGT GTA G GGA AAG AGT GTA GAT CTC GGT GGT CGC /SpacerC18/ CAC TCA /SpacerC18/ TTC AGA CGT GTG CTC TTC CGA TCT ATT GAT GGT GCC TAC AG), followed by circularization with Circligase (Epicenter, USA). Circularized cDNA was then PCR amplified using primer NINI2 (AAT GAT ACG GCG ACC ACC GAG ATC TAC AC) and a primer with a barcode (CAA GCA GAA GAC GGC ATA CGA GAT XXX XXX GTG ACT GGA GTT CAG ACG TGT GCT CTT CCG), where XXXXXX denotes a six-nucleotide-barcode used to distinguish samples run in the same lane (Table S4). For RNA-seq, total RNA was isolated from the same cell extracts using SDS/hot acid phenol/chloroform extraction. The Ribo-Zero Gold Yeast kit was used to remove rRNA, and total RNA was randomly fragmented by incubating for 20 minutes at 95°C in freshly made fragmentation buffer (100 mM sodium carbonate-bicarbonate [pH 9.2], 2 mM EDTA). RNA was then precipitated and fragments of 40–60 nt were purified from a denaturing PAGE gel, and library generation carried out as above. Ribo-Seq and RNA-Seq were performed for two independent cultures for each condition (WT and Δntd cells grown in SD both with and without 3% Urea), and the 16 libraries sequenced in 2 lanes with 150 bp reads on an Illumina HiSeq 4000 instrument by Genewiz.

### Analysis of ribosome profiling libraries

The Ribogalaxy platform (https://ribogalaxy.ucc.ie, (30)) was used for trimming linker sequences (Cutadapt version 1.1.a; (31)), subtractive alignment of *S. cerevisiae* non-coding RNAs (Bowtie version 1.1.a; (32) using the R64.2.1 S288C genome from Saccharomyces Genome Database (SGD, RefSeq ID: 285498), alignment of rRNA subtracted libraries to the transcriptome (Bowtie version 0.1.3 using the SGD transcriptome dataset and counting of uniquely mapped reads (27) using Ribocount version 0.3.1. Statistical analyses of differences in total RNA counts, ribosome footprints, or TE values between WT, mutant, urea-treated, and untreated samples were conducted using DESeq2 (33). Gene ontology categorization of library-specific differences was performed at SGD.

### qRT-PCR

Cell lysates were prepared and 12 fractions were collected from polysomes fractionated using the protocol for riboseq gradient preparation above (without nuclease treatment.) 0.3 ml of each gradient fraction was spiked with equal amounts of control RNA (Fluc mRNA, Trilink Biotechnologies, USA), then total RNA was extracted using the hot acid phenol-chloroform method (34). First strand synthesis was performed with iScript Advanced Reverse Transcriptase (Biorad, USA) using oligo-dT primers and random hexamer primers. Quantitative PCR was performed with iQ SYBR Supermix reagents (Biorad, USA) using CFX384 Touch Real-Time PCR detection system (Biorad, USA) two times per sample. Gene-specific primer sequences are listed in supplementary table 3.

## RESULTS

### The N-terminal domain of eIF4B increases the ratio of polysomes to monosomes in yeast

Previous work demonstrated that the N-terminal domain (NTD) of eIF4B promotes both affinity for the 40S subunit in vitro and recruitment of mRNAs to the preinitiation complex in vitro and in vivo, while the RRM of eIF4B is dispensable in auxotrophic yeast strains (13,17). Given the diverse dependencies of cellular mRNAs on eIF4B function (6), we thought it likely that differential mRNA selection promoted by eIF4B could confer phenotypic advantages. We wondered if the ability of the NTD to promote translation would afford cells the ability to resist different stressors, and whether the RRM could provide additional function under stress, so we constructed prototrophic strains for phenotype microarray analysis. We previously reported that deletion of the NTD conferred slow growth and cold-sensitivity in a strain auxotrophic for histidine, uracil, and methionine on solid media (7,13). For this work, an eIF4B null mutant was transformed with a plasmid that complemented all four existing auxotrophic markers and provided a WT, Δntd, or Δrrm eIF4B gene copy (*TIF3, tif3 Δntd*, or *tif3* Δ*rrm*) under the native promoter and terminator (Fig. 1A). As previously reported, deletion of the NTD, but not the RRM reduced growth rate (Fig. 2B,(13)).

Polysome profiles from these prototrophic strains expressing WT, Δntd, and Δrrm eIF4B confirmed that NTD deletion led to a gross reduction of polysomes and an increase in monosomes in the mutant (Fig. 1B, S2), indicating the NTD promotes global translation initiation in vivo in a prototrophic background. RRM deletion had only a minor effect on polysome to monosome ratio when expressed from this single copy plasmid (Fig. 1B, blue), in agreement with our previous findings (13).

### Yeast eIF4B associates with ribosome complexes in vivo

We previously reported that deletion of the NTD decreased binding affinity of eIF4B for purified 40S subunits. In addition, deletion of the NTD decreased the rate constants and endpoints of mRNA binding to the PIC in vitro, while affecting the conformation of two areas of the rRNA near protein RPS20/uS10 (13). To determine whether deletion of the NTD affects association of eIF4B with ribosomes in yeast, we performed velocity gradient fractionation of formaldehyde-crosslinked lysates followed by Western blotting of eIF4B, eIF4A, eIF4G, and RPS2. We performed two types of gradients to observe changes in association of eIF4B with both translating ribosome complexes and 40S subunits and PICs as a function of NTD and RRM deletion (Fig. 1C-F, Fig. S1). Running crosslinked lysates on a 5-45% gradient effectively separates polysome-, monosome-, and mRNP-fractions. Importantly, when quantifying the fraction of eIF4B in each fraction of the gradient by Western blotting, we found that ~55% of WT eIF4B comigrated with Rps2/40S subunit-containing fractions (Fig.1C-D fractions 4-10), both as part of the 40S/PIC and moreso with the later fractions containing translating polysome complexes, which make up more of the ribosome pool. Upon deletion of the NTD we saw that Rps2 shifted from later to earlier fractions, confirming the polysome to monosome shift observed by UV spectroscopy (Compare Figure 1 panels B and D). This indicates deletion of the NTD led to less ribosomes associated with mRNAs, suggesting reduced translation initiation rate in these cells. In addition, we found that upon deletion of the NTD, eIF4B position in the gradient was shifted such that ~80% of the protein moved to the first two fractions that lack 40S subunits and ribosomes (Fig. 1C, compare black and red.) This is in contrast to deletion of the RRM, which conferred only a minor decrease in translating ribosome capacity, judged by similar polysome:monosome ratio as WT (Fig. 1B, compare black and blue), and also did not grossly affect eIF4B association with translating ribosomes (Fig. 1C, D.)

Ultracentrifugation of crosslinked lysates on 7.5-30% sucrose gradients optimally separates 40S/PIC fractions (Fig. S1.) We found that WT eIF4B was present in fractions 10-13 that contain RPS2 and indicate 40S, 43S, and 48S complexes. As we saw in Figure 1, eIF4B was also present in early fractions containing mRNPs (Figure S1A, C). Deletion of the NTD reduced the amount of eIF4B in these Rps2-containing fractions by 93%, with increased eIF4B in early fractions containing proteins and mRNPs.

We also found that while the RRM did not change the amount of eIF4B comigrating with the overall ribosome pool (Figure 1), deletion of this domain did decrease occupancy of eIF4B on the fraction of the 40S subunits present in PICs, although not to the extent observed for NTD deletion, which nearly eliminated eIF4B occupancy in PIC fractions (Figure S1). It is notable that deletion of the RRM decreased the concentration of eIF4B in cells (Figure 2D), and was previously shown to have a minor effect on 40S binding affinity and apparent affinity for the PIC in an mRNA recruitment assay (13), so this decrease in RRM occupancy of PICs outside of polysomes could be a reflection of that decreased ribosome affinity.

As a control we blotted the same gradient fractions for additional 48S components, eIF4A and/or eIF4G (Fig. 1E, F and Fig. S1B). In contrast to eIF4B, we found that eIF4A and eIF4G remained distributed across gradient fractions when the NTD or RRM of eIF4B were deleted. This suggests the affinity of eIF4A and eIF4G for the PIC are not mediated by eIF4B. Moreover, this suggests that shift of Δntd eIF4B from polysome and 40S fractions to ribosome-free fractions is the result of decreased ribosome binding affinity in the mutant, since these other components of 48S PICs did not show a PIC and polysome to mRNP shift.

### The NTD of eIF4B promotes translation-mediated stress responses

To determine conditions under which the NTD of eIF4B plays a specific role in regulating translation, we performed phenotype microarrays of cells expressing WT or Δntd eIF4B from a single copy plasmid that also restored nutrient prototrophy. Phenotype microarray analysis showed a large number of conditions in which the prototrophic Δntd eIF4B expressing strain grew at a reduced rate compared to the wild-type eIF4B expressing cells (Figure 2A). The strongest responses (Table S5) include osmolytes, detergents, a number of peptides as nitrogen sources, and antibiotics. Urea, which gave the strongest negative phenotype when present at 3% w/v in the media, acts as a denaturing agent and can cause membrane blebbing at high concentrations (35,36), and can readily cross the yeast cell wall and membrane to act as a nitrogen source (37). Tamoxifen, which targets the estrogen receptor in higher eukaryotes, targets the calmodulin protein in yeast which regulates stress responses through Hog1 interaction (38,39). Poly-L-Lysine can act as a cationic detergent or a charged adherent for various molecules, and likely interacts with the cell wall or membrane. The strain lacking the NTD of eIF4B also showed heightened sensitivity to antibiotics that target the small ribosomal subunit, apramycin sulfate and to a lesser extent tobramycin. WT yeast are not sensitive to these antibiotics, which when combined with the sensitivity to various salts (Potassium chloride, Chromium chloride, and to a lesser extent, sodium chloride) and other phenotypes described above, suggests a defect in membrane and/or cell wall permeability in the mutant. Growth defects were verified for Δntd-expressing cells in the presence of urea, which conferred the largest reduction in the mutant (Figure 2B, red). These experiments confirmed that WT growth rate is unaffected by urea (Figure 2B, black) while the mutant shows slow growth. In contrast, deletion of the RNA-binding RRM domain, which diminishes in vitro RNA-binding affinity(13), conferred no large reproducible advantage or hindrance, suggesting the RNA-binding activity of eIF4B, at least by the RRM, is dispensable for growth in all conditions tested (Figure 2B, blue). Together these results suggest the eIF4B 40S-binding NTD allows resistance to a number of growth conditions that challenge cellular integrity, and that the mutant may have a defect in membrane and/or cell wall permeability.

To further investigate the mechanisms by which the NTD promotes growth in the presence of stressors, we compared polysome traces for WT and Δntd eIF4B-expressing cells grown in the presence and absence of 2% or 3% urea (Figure 2C, traces shown in Figure S2). Whereas WT eIF4B-containing cells showed minor decreases in polysome to monosome (P:M) ratio upon addition of either concentration of urea (7% reduction in 2% and 10% reduction in 3% urea; Figure 2C, black), Δntd cells showed further 36% and 50% decreases in P:M ratio due to urea exposure (Figure 2C, red). In contrast, deletion of the RRM resulted in less than 5% change in P:M ratio (Figure 2C, blue).

The reduction in translation conferred by NTD deletion is not due to altered levels of eIF4B, as immunoblotting for a His tag on the eIF4B C-terminus shows similar protein levels when grown in media with varied urea concentrations (Figure 2D). As noted before, deletion of the RRM resulted in reduced detection of that protein. However, the level of detectable protein was unaffected by urea addition. Together these data suggest interactions and activities promoted by the NTD allow robust translation in the presence of urea.

### The eIF4B NTD promotes translation of mRNAs encoding membrane-associated proteins

We next performed ribosome profiling on the yeast cells expressing WT or Δntd eIF4B, both with and without urea to determine the mechanism by which eIF4B·40S leads to changes in translation of individual mRNAs. Ribosome profiling maps the positions of translating ribosomes on mRNAs to determine which sequences are translated more and less effectively in response to changes. We prepared illumina-indexed cDNA libraries for RNAseq and Riboseq from WT and Δntd eIF4B-expressing cells in the presence or absence of 3% urea. Comparison of replicates indicates sufficient reproducibility for each sample, with slightly higher variability in the Riboseq libraries from Δntd eIF4B-expressing cells with 3% urea (Figure S3), which showed the strongest global repression of translation and therefore had the least ribosome footprints. After mapping and quantifying footprints on mRNAs in the yeast transcriptome, we compared the averaged changes in RNAseq, Riboseq, and TE (translation efficiency, Riboseq/RNAseq) in response to urea exposure in WT versus the urea-dependent change in Δntd eIF4B-expressing cells (Figure 3, Table S6). Changes in individual mRNA levels in response to urea were more similar between the two strains (Figure 3A, r=0.622), while changes in ribosome footprints and changes in TE for individual mRNAs in response to urea showed less correlation between the two strains (Figure 3B, Riboseq, r=0.361; 3C, TE, r=0.156). This is further evidenced by evaluation of the mRNAs with ≥ 1.5-fold increase in RNAseq, Riboseq and TE in response to urea. There were fewer mRNAs overall showing ≥1.5-fold urea-dependent increases in RNA levels (Figure 3D; WT= 39, Δntd=108) and substantial overlap between the WT and mutant pools (43.5% of mRNAs showing ≥1.5-fold increase for WT eIF4B cells also increased by ≥1.5-fold in Δntd). In contrast, there were large numbers of mRNAs with ≥1.5-fold changes in RiboSeq (Figure 3E, WT=141 mRNAs, Δntd=543 mRNAs) and TE (Figure 3F, WT=300 mRNAs, Δntd=163 mRNAs), both in the WT and Δntd eIF4B-expressing cells. Moreover, the pools of mRNAs showing these changes in TE upon exposure to urea were almost completely distinct in the control cells versus those mRNAs whose translation increased in the Δntd mutant. It is important to note that while there are a greater number of individual mRNAs showing a relative increase in Riboseq in the mutant in response to urea, these cells show an overall decrease in global translation initiation capacity as evidenced by decreased P:M ratio (Figure 1B, 2C).

**Figure 3.**
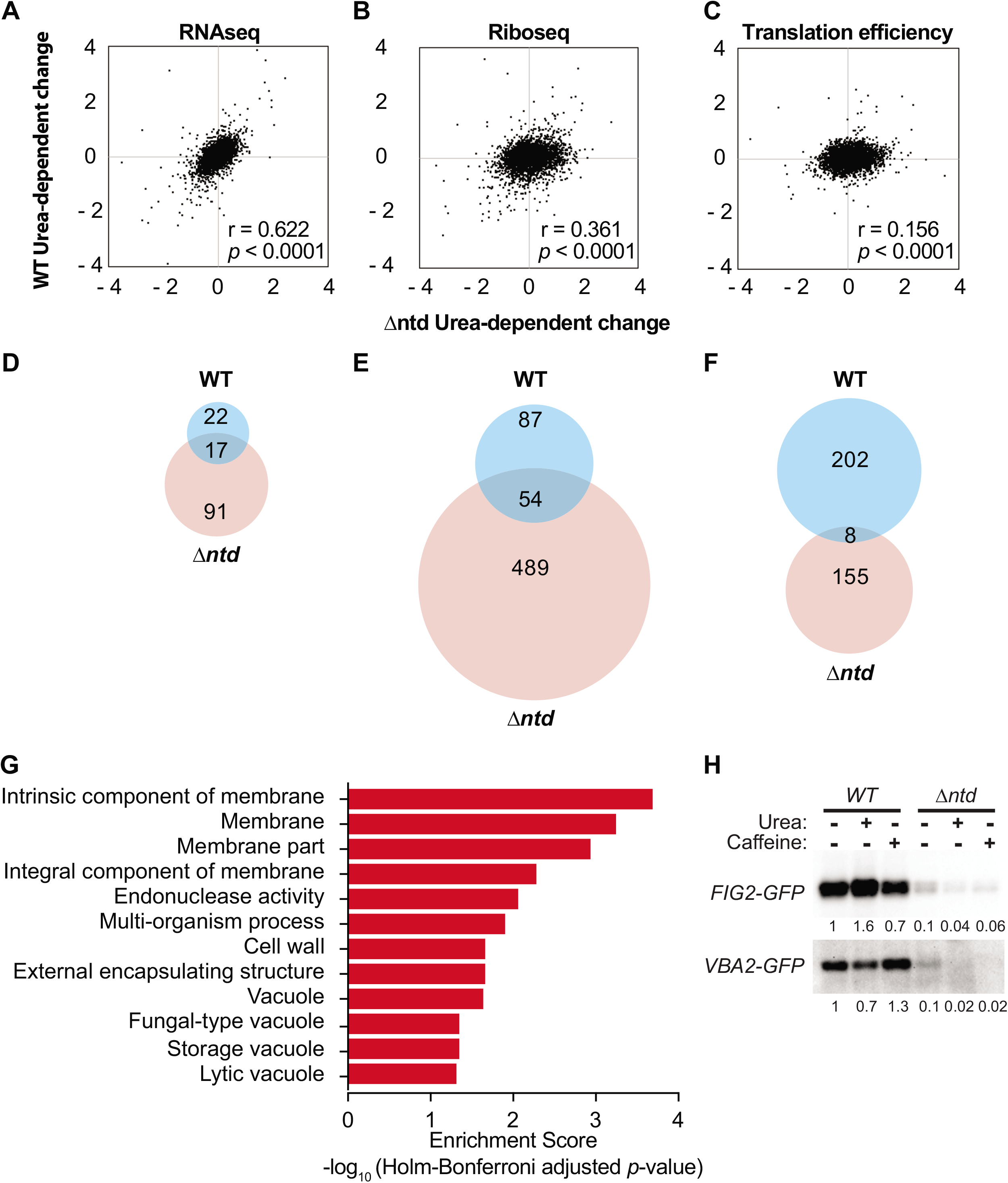
The NTD of eIF4B promotes translation of mRNAs encoding proteins associated with the membrane and cell wall. **A-C**. Comparison of Log2 values for changes in RNAseq (**A**), Riboseq (**B)**, or TE (**C**, Translation efficiency) in response to 3% urea for WT are plotted vs. the Log2 values for changes observed in the NTD deletion mutant for each of the 4070 genes with measurable expression in each group. Pearson correlation coefficients are shown. **D-F**. Overlap of urea-dependent genes exhibiting 1.5-fold or greater increase in RNAseq (**D**), Riboseq (**E**), or TE (**F**) for WT and Δ*ntd*. **G**. Gene ontology analysis for mRNAs with increased TE in WT in response to urea. **H.** Western analysis of GFP translation reporters. The 5’UTR and first 30 nucleotides of the FIG2 and VBA2 genes were fused to GFP in a plasmid under the native promoters for each. Indicated transformants of WT and Δntd eIF4B-expressing yeast were subjected to anti-GFP western analysis following growth in the absence or presence of 3% urea or 1.5 mg/ml caffeine. The fractions of reporter band intensity per total protein bands for each lane on the gel were normalized to WT eIF4B without additive for each reporter.

Changes in ribosome occupancy of representative mRNAs were verified by performing qRT-PCR on gradient fractions from cells expressing WT and Δntd eIF4B in the presence and absence of 3% urea in the growth media. The trends observed by ribosome profiling are confirmed by this independent method for several representative mRNAs in each pool (Figure S4). Moreover, to determine whether changes in ribosome occupancy correlated with changes in protein production, we designed two translation reporters in which the 5’ UTR and first 30 nucleotides of two mRNAs that showed NTD-dependent reductions (FIG2 and VBA2) were cloned in frame in front of a GFP gene (Table S2). Western blotting for GFP in extracts of cells harboring both a FIG2-GFP reporter and WT or Δntd eIF4B showed that addition of urea to WT cultures increased the level of FIG2-GFP protein by 1.6-fold, while addition of caffeine slightly decreased FIG2 production (relative to total protein per lane on gel) (Figure 3H). In contrast, addition of urea to WT cells did not increase steady state levels of VBA2-GFP, while addition of caffeine slightly increased production by 1.3-fold (Figure 3H). This result is consistent with the increased polysome association observed for FIG2 upon urea addition, but little to no change in polysome association observed for VBA2 in WT cells (Fig. S4). Importantly, deletion of the NTD reduced the steady state levels of both reporter proteins by 90% or more, supporting the conclusion that reduced ribosome association upon NTD deletion led to reduced protein production from these mRNAs. Together these data indicate that the eIF4B·40S binding NTD promotes changes in growth by differentially affecting translation of specific mRNAs in response to stressors.

To understand how eIF4B NTD-dependent changes in translation promote growth in urea, we first performed gene ontology (GO) analysis of the mRNAs showing ≥1.5-fold increased TE in response to urea in WT or mutant cells (Figure 3G, WT). Of the 210 mRNAs with ≥1.5-fold increased translation in WT cells upon exposure to urea, 102 mRNAs were associated with the parental membrane (GO) term, while significant numbers of mRNAs were associated with the cell wall, cellular periphery, and other related terms, suggesting the eIF4B NTD enhanced translation of mRNAs encoding proteins that remodel or otherwise localize to the cellular membrane. In contrast, of 163 mRNAs showing ≥1.5-fold increased translation in Δntd-expressing cells in response to urea, 151 of those mRNAs were associated with the cytoplasm gene ontology term (Table S7). Likewise, analysis of the mRNAs showing increased TE in Δntd cells showed strong association with GO terms for ribosomes and cytosolic components, even without urea (Table S8). Furthermore, mRNAs that showed decreased translation when the eIF4B NTD was removed in the absence of urea were associated with membrane-bounded organelles (Table S9). This suggests the NTD promotes translation of mRNAs encoding membrane-associated proteins, and the loss of this ability results in the mutant translating mRNAs that encode cytoplasmic proteins, leading to urea and other stress sensitivities. Finally, we analysed the 281 mRNAs showing more than 1.5-fold decreased translation efficiency in response to urea in the mutant cells relative to WT. In this case we saw decreased translation for 84 mRNAs associated with the endomembrane system (p-value=0.000026), including association with the ER, Golgi, transferase activities, glycosylation, and mannosylation (Table 1). This suggests effective translation of mRNAs for proteins trafficked through the ER/Golgi network to the membrane and cell wall is dependent on the eIF4B NTD activities. Interestingly, the specific mRNAs that were dependent on the eIF4B NTD showed little overlap with those mRNAs identified as hyperdependent on eIF4B when analysing the translatome of an eIF4B null mutant (data not shown, (6)) despite sharing general structural features discussed below. However, in the previous analysis of cells lacking eIF4B, cells were shifted to 37°C for 2 hours prior to ribosome profiling, so differences in translation of individual mRNAs could also be due to this variable. Together these results suggest that the eIF4B 40S-binding NTD is necessary for critical changes in translation of RNAs that remodel the cellular periphery in response to urea exposure. This function is necessary for translation of the optimal pool of mRNAs to promote rapid growth in standard media as well.

**Table 1:** Gene Ontology analysis for genes with decreased translational efficiency in Δ*ntd* compared to WT in the presence of urea

| GO Term | p-Value |
| --- | --- |
| Transferase activity, transferring hexosyl groups [GO:0016758] | 1.62E-06 |
| Endomembrane system [GO:0012505] | 1.44E-05 |
| Transferase activity, transferring glycosyl groups [GO:0016757] | 2.22E-04 |
| Protein glycosylation [GO:0006486] | 3.76E-04 |
| Macromolecule glycosylation [GO:0043413] | 3.76E-04 |
| Mannosylation [GO:0097502] | 5.19E-04 |
| Glycoprotein biosynthetic process [GO:0009101] | 8.19E-04 |
| Cellular bud [GO:0005933] | 1.55E-03 |
| Protein N-linked glycosylation [GO:0006487] | 1.91E-03 |
| Glycosylation [GO:0070085] | 2.40E-03 |
| Glycoprotein metabolic process [GO:0009100] | 2.85E-03 |
| Cell periphery [GO:0071944] | 4.09E-03 |
| Golgi cisterna [GO:0031985] | 4.70E-03 |
| Cell part [GO:0044464] | 8.14E-03 |
| Cell [GO:0005623] | 8.81E-03 |
| Protein O-linked glycosylation [GO:0006493] | 1.98E-02 |
| Golgi stack [GO:0005795] | 2.26E-02 |
| Mannosyltransferase activity [GO:0000030] | 3.04E-02 |
| Organelle subcompartment [GO:0031984] | 3.65E-02 |
| Membrane part [GO:0044425] | 4.69E-02 |

### The NTD of eIF4B promotes translation of mRNAs with long and structured 5’UTRs

Previous studies have shown that mammalian eIF4B is necessary for PIC assembly at the start codon of mRNAs with 5’UTR secondary structure in vitro (4). Likewise, yeast eIF4B is associated with robust loading and scanning of PICs and translation of mRNAs and synthetic reporters containing higher than average secondary structure in vitro and in vivo (5,6). We speculated that the effect of the eIF4B NTD on translation of mRNAs needed to combat extracellular urea is related to the ability of eIF4B to promote translation of structured mRNAs. To test this hypothesis, we analysed available parallel analysis of RNA structure (PARS) scores (40) for the mRNAs exhibiting changes in translation as a result of eIF4B NTD deletion, in the presence or absence of urea. PARS scores provide the relative propensity of each nucleotide to a single- or double-strand specific nuclease, with a higher PARS score indicating a higher propensity for secondary structure. Cumulative PARS scores can be compared for specific regions of mRNAs to determine the likelihood that the e.g. first 30 nucleotides, total 5’ UTR, or regions in the ORF have more structure (Figure 4A), which would present an impediment for PIC loading, PIC scanning, or translation elongation respectively (6). We found that deletion of eIF4B led to lower translation efficiency (≥1.5-fold decrease) of mRNAs with significantly higher PARS scores for the first 30 nucleotides of the 5’UTR (Figure 4B, First30). An even larger difference was observed for the total 5’UTR, suggesting interactions of the eIF4B NTD promote effective mRNA loading and possibly scanning through structured mRNA 5’UTRs (Figure 4B, 5’UTR). The average individual nucleotide PARS score averaged for the full 5’UTRs was likewise significantly higher for these groups of mRNAs that were hyperdependent on the eIF4B NTD for translation (Figure 4C). In contrast, there was not a significant change in the PARS scores for the 30 nucleotides surrounding the start codon (Figure 4B, Start30), or the first 30 nucleotides of the ORF, suggesting structure around the start site and in the ORF does not strongly require eIF4B activity. In fact, the PARS scores for the Plus45, Plus60 and Plus75 regions were significantly lower for the group of mRNAs that were hyperdependent on the NTD of eIF4B (in the absence of urea). This suggests that the eIF4B NTD is not required for translation of mRNAs with structured ORFs or structured RNAs in general, but instead is important for PIC loading and movement through structured 5’UTRs (Figure 4A, B). The mean length of 5’UTR was also significantly higher for the group of mRNAs that were less efficiently translated (≥1.5-fold) when the NTD of eIF4B was deleted. This further suggests the NTD is critical for effective scanning of long structured 5’UTRs (Figure 4D). This effect was more pronounced when cells were grown in the presence of 3% urea prior to ribosome profiling, suggesting the effect of the NTD on urea resistance stems from the ability of eIF4B to promote effective scanning.

**Figure 4.**
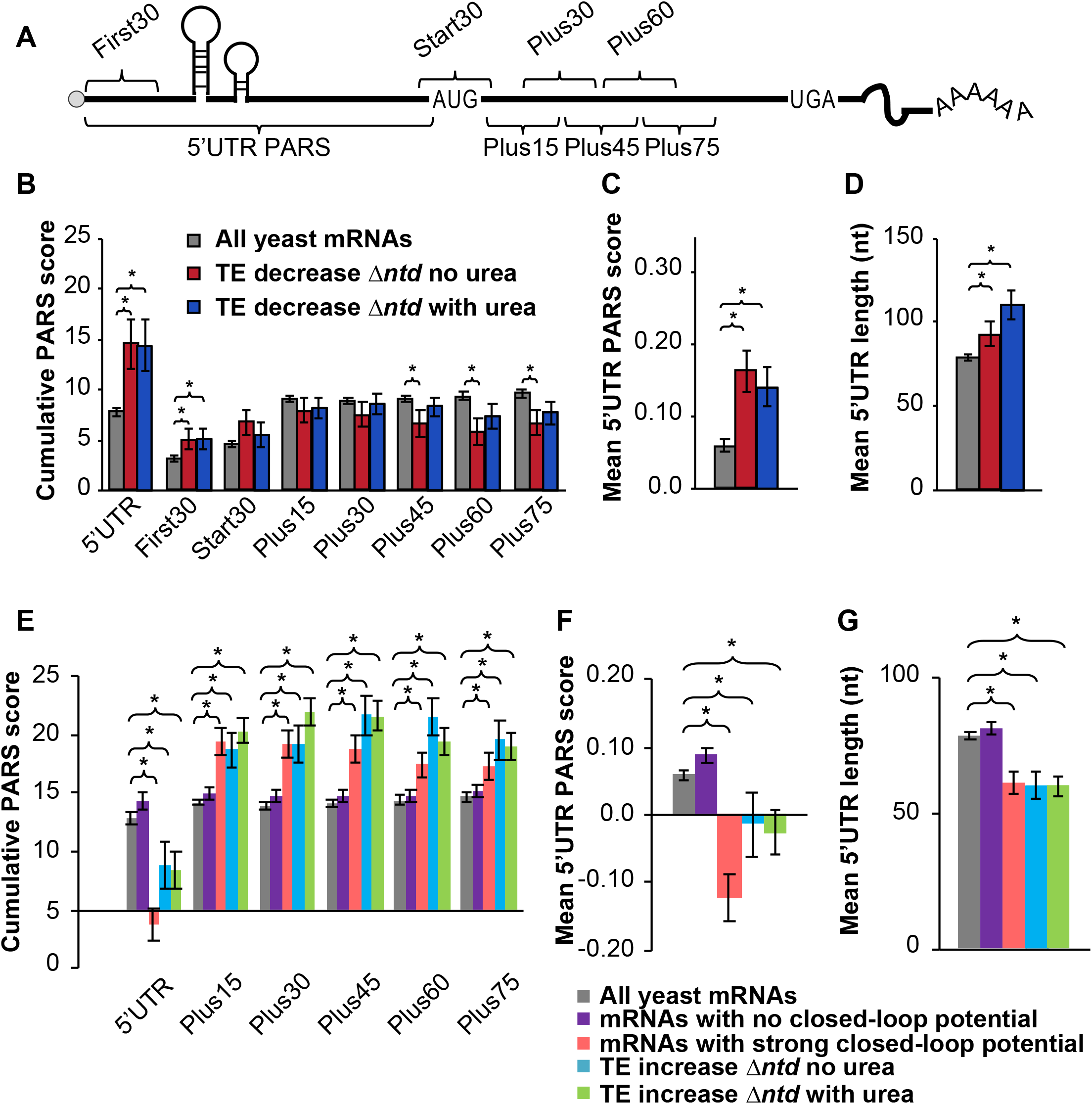
Comparison of PARS (Parallel analysis of RNA structure) scores indicates a higher propensity for secondary structure in the 5’UTRs of mRNAs that are dependent on the NTD for translation, and less associated with closed-loop factors. **A.** Schematic showing 5’-UTR and CDS intervals for cumulative PARS scores. The sum of scores for all 5’-UTR nucleotides (5’UTR PARS); for the first 30 nucleotides (First 30 PARS); for 30 nucleotides surrounding the start codon (Start 30 PARS); and for nucleotides within the ORF, from +1 to +30 (Plus15), +16 to +45 (Plus30), +31 to +60 (Plus45), +46 to +75 (Plus60), and +61 to +90 (Plus75). **B-C.** The mean PARS scores (calculated from data reported in reference (40)) for indicated cumulative regions (**B**), or for individual nucleotides in the 5’UTR (**C**) are indicated for all yeast mRNAs with available PARS scores (gray, n=2679); mRNAs with decreased TE (≥1.5-fold) in Δ*ntd* relative to WT in media lacking urea (red, n=138); and for mRNAs with decreased TE in Δ*ntd* relative to WT in 3% urea (navy, n=156). **D**. Average length of 5’-UTR for the indicated sets of mRNAs. **E-G**. PARS and 5’UTR length analysis calculated for the indicated gene sets, with P values from Student’s t test indicated (*P < 0.05): grey bar, all yeast mRNAs with available PARS scores (n = 2679); purple bar, mRNAs with no closed-loop potential, characterized for de-enrichment in immunoprecipitation of eIF4F and Pab1, and enrichment in immunoprecipitation of eIF4E-binding proteins as shown in (41); red bar, mRNAs with strong closed-loop potential, characterized for de-enrichment in immunoprecipitation of eIF4E-binding proteins and enrichment in immunoprecipitation of eIF4F and Pab1, as shown in (41); blue bar, mRNAs with increased TE in NTD deletion mutant as compared to WT without urea; green bar, mRNAs with increased TE in NTD deletion mutant as compared to WT in 3% urea. **E.** Average PARS scores calculated for the indicated sets of mRNAs for each 5’-UTR or CDS interval described in Figure 4A. **F.** Average PARS score calculated for entire 5’-UTR for the indicated sets of mRNAs. **G.** Average length of 5’-UTR for the indicated sets of mRNAs.

We also found that 155 mRNAs showed a relative increase in translation in Δntd cells (Figure 3F), indicating ribosomes were able to be loaded on these mRNAs without eIF4B NTD activities. We assessed the degree of secondary structure in the 5’UTRs and coding sequences of these mRNAs (Figure 4E-G, cyan and green). Previous work investigated mRNAs translated relatively more efficiently in a mutant lacking eIF4B, and noted that these mRNAs also displayed increased association with components of the closed loop complex (6): eIF4E, eIF4G and PABP (41). We compared mRNAs classified as strong-closed loop potential (higher crosslinking immunoprecipitation association with closed-loop components) and no closed-loop (enriched in inhibitors of the closed-loop complex(41)) to mRNAs that were translated ≥1.5-fold more efficiently in the Δ*ntd* mutant. We found that mRNAs that showed increased translation in Δntd (with or without urea, cyan and green) showed similar trends with respect to PARS scores as those mRNAs defined as having strong closed-loop potential, and the opposite behavior as those mRNAs defined as having no closed-loop potential. Both the Total 5’UTR region and the average per nucleotide PARS scores for the 5’UTRs of these mRNAs showing eIF4B NTD-independence were significantly lower than the average yeast mRNA. In contrast, the mRNAs that showed increased translation efficiency in the Δntd mutant and strong-closed-loop associated mRNAs were more structured than the average yeast mRNA in the ORF. This suggests that mRNAs that rely on closed-loop components for mRNA loading do not require the eIF4B NTD or its interaction with the ribosome (13), and reinforces the conclusions of the previous manuscript that mRNAs requiring eIF4B activity are less associated with closed-loop components (6).

### RNAs encoding proteins trafficked through the ER and Golgi have long and structured 5’UTRs, imposing a heightened requirement for eIF4B

Phenotype microarray analysis suggested the NTD of eIF4B stimulated growth in a number of diverse conditions that challenge cellular integrity (Figure 2A, Table S5). The findings that mRNAs translated more effectively by full-length eIF4B had longer and more structured 5’-untranslated regions than those translated when the NTD was deleted led us to question whether proteins for different functions in cells may rely on distinct translational mechanisms. For instance, mRNAs for proteins that promote rapid growth may have less structure and rely less on eIF4B, whereas proteins that allow adaption to stressors, such as the membrane and cell wall proteins, may have more structure and require eIF4B function for translation. If true, the degree of structure would be expected to impose regulatory capacity as cells encounter stresses that require membrane changes, and may explain how the NTD of eIF4B affords resistance to diverse stressors that may require different membrane composition. To investigate this further, we compared the PARS scores for all yeast mRNAs versus the PARS scores for all yeast mRNAs associated with GO terms for mRNAs that required eIF4B for translation in response to urea (Figure 5A-C.) This group includes: intrinsic component of the membrane (the GO term with the lowest P value for RNAs showing increased translation in response to urea in WT cells); as well as transferase activity, endomembrane system, glycosylation, and mannosylation (parent GO terms for mRNAs with decreased translation in response to urea in Δ*ntd* cells.) Interestingly, we found that the 5’UTRs of mRNAs associated with each of these GO terms had higher average Total PARS scores than the average yeast mRNA (Figure 5A, 5’UTR). However, only those mRNAs encoding intrinsic components of the membrane had significantly longer 5’UTRs (Figure 5B). The mean 5’UTR PARS scores for individual nucleotides was also significantly higher than the average yeast mRNA for all classes (Figure 5C), indicating these classes of mRNAs associated with dependence on the ribosome binding NTD of eIF4B have inherently more structure in the 5’UTRs. This suggests functional importance of structural elements in regulating translation of membrane-associated and trafficked proteins. We then compared the structural composition of mRNAs from two gene ontology categories that were enriched under eIF4B NTD independent translation (Figure 5D-F). We found that as expected, given the observed NTD-independent translation associated with these classes of mRNAs, cytoplasmic translation and structural constituent of the ribosome mRNA categories as whole showed a dearth of structure in their 5’UTRs, with overall negative cumulative PARS scores and mean 5’UTR scores per nucleotide, indicating the 5’UTR regions of these mRNAs are likely to be single stranded. Interestingly, the cytoplasmic translation mRNAs had no significant difference in lengths of their 5’UTRs from the pool of all yeast mRNAs. In contrast to the 5’UTR however, the regions just downstream of the start codon showed higher than average PARS scores for these translation-associated gene ontology classes (Figure 5D, blue and red).

**Figure 5.**
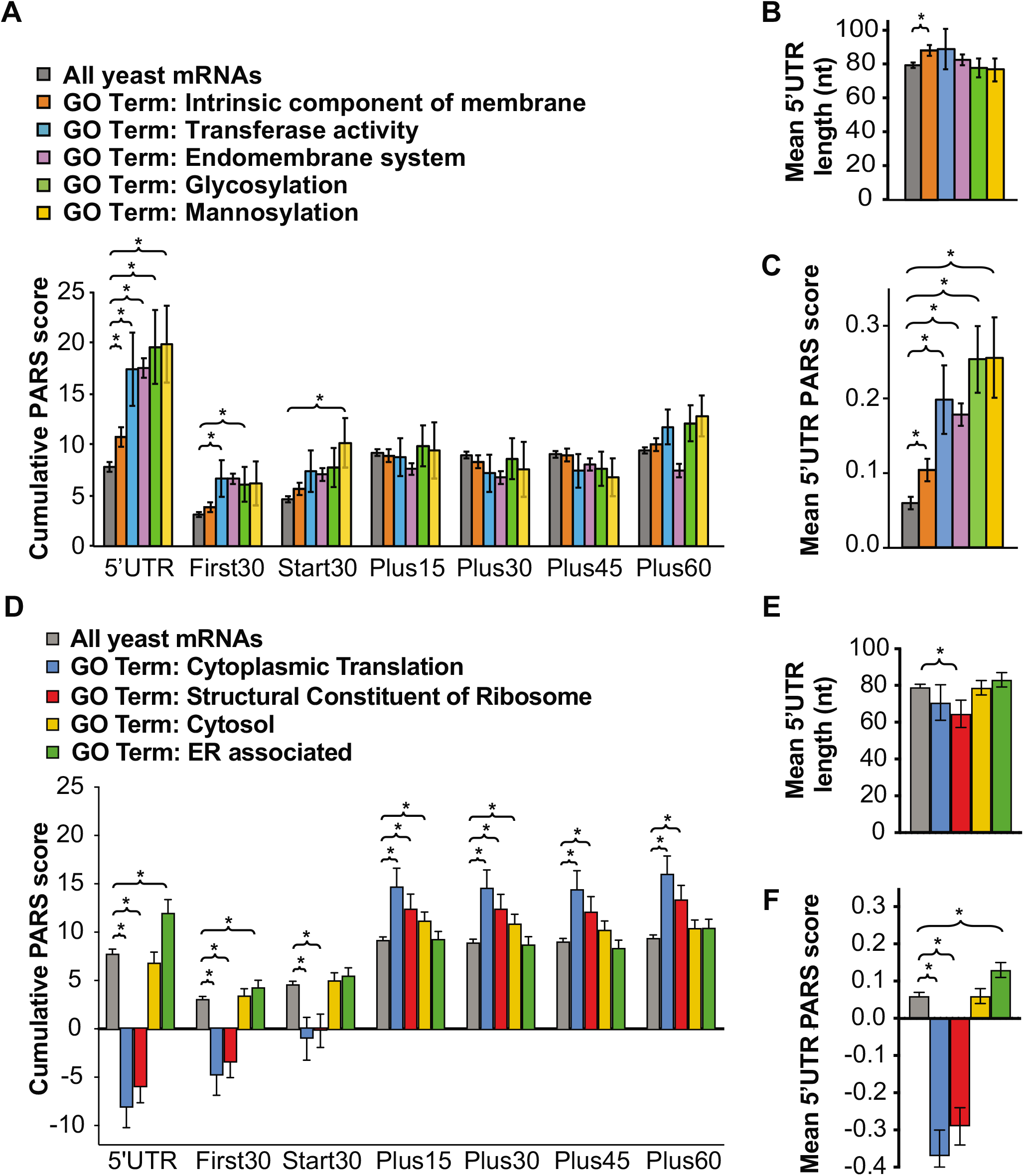
RNAs encoding proteins trafficked through the ER and Golgi have long and structured 5’UTRs, imposing a heightened requirement for eIF4B. Averaged cumulative (A,D) and average 5’UTR nucleotide (C, F) PARS scores and 5’UTR lengths (B, E) for all genes associated with indicated gene ontology categories: intrinsic component of membrane (orange, n=1360), transferase (A-C, blue, n=86), endomembrane system (purple, n=1098), glycosylation (A-C, green, n=87), mannosylation (AC, yellow, n=46), cytoplasmic translation (D-F, blue, n=161), structural constituent of ribosome (red, n=190), cytosol (D-F, yellow n=426), or ER associated (D-F, green, n=405) for each 5’-UTR or CDS interval described as in Figure 4A, with P values from Student’s t test indicated (*P < 0.05)

We finally compared the structural content of the two broad gene ontology classes: cytosol and ER. Whereas the cytosol class showed no significant difference in 5’UTR PARs scores from average yeast mRNA, the ER-associated gene ontology class showed significantly higher Total 5’UTR and mean 5’UTR PARS scores than all yeast mRNAs. Moreover, the open reading frames of the ER-associated mRNA pool showed the opposite trend. The cytosol class showed slightly elevated PARS scores for the region immediately downstream of the start site than observed for all yeast mRNAs. Together this suggests higher structure in the 5’UTRs of ER-associated mRNAs than cytosolic mRNAs.

We also took an unbiased approach to exploring the relationship between gene ontology classes, 5’UTR features, and eIF4B NTD-dependence. We ranked all yeast mRNAs based on their cumulative 5’UTR PARS scores (Figure S5A-C) or 5’UTR lengths (Figure S5D-F) and performed gene ontology analysis to determine enrichment of specific biological processes for the top (B, E) and bottom (C, F) 30% of mRNAs from each group. We compared the degree of overlap between the resulting GO term lists, and found that eIF4B NTD-independence, low 5’UTR structure propensity, and short 5’UTR gene ontology terms showed striking overlap, particularly for the highest enriched GO terms. These mRNAs encode proteins associated with cytoplasmic translation, ribosome biogenesis, and other processes related to ramping up protein synthesis. In contrast, those GO terms enriched for transcripts exhibiting higher eIF4B NTD-dependence showed some overlap with those enriched in mRNAs with high 5’UTR structure, but considerably less overlap with those enriched for mRNAs with long 5’UTRs. Overall, these data suggest a complex relationship between the ability of eIF4B to promote translation of mRNAs with structured 5’UTRs and regulation of translation that promotes growth versus regulatory changes.

## DISCUSSION

In this study, we characterized the contribution of eIF4B RNA- and 40S subunit-binding domains to translational control as well as the ability to promote adaptation of yeast to diverse stressors. We found that the NTD of eIF4B promoted association of eIF4B with PICs and polysomes in yeast while allowing translation of mRNAs with longer than average and highly structured 5’UTRs.The NTD also afforded yeast the ability to translate mRNAs encoding proteins trafficked through and modified in the ER and Golgi to reside in cellular membranes. These proteins are expected to remodel the cellular periphery and allow yeast to cope with external stressors.

The RRM of eIF4B was thought to promote mRNA recruitment to ribosomes by providing an RNA anchoring point on a ribosome or eIF3-bound molecule (namely for mammalian eIF4B, (42–45)) or by promoting RNA strand-exchange activities of eIF4B (16). Our previous work suggested that instead, the RNA-binding activities of the RRM are dispensable for eIF4B function in yeast (13). However, because the experiments in our previous work analysed the function of Δrrm-expressing eIF4B under optimal growth conditions, it remained plausible that the RRM provides additional functions to cells under stress, when additional interactions may be needed to direct ribosomes to specific mRNAs. Our phenotype microarray analysis of the Δ*rrm* mutant provides strong evidence that the RRM domain is in fact dispensable for function of this protein in yeast, at least in liquid media. The only plate in which we saw mild phenotypes for the Δ*rrm* mutant was in the presence of certain alternative sulfur sources (Figure 1A), but the changes observed were well below the cutoff for significance and were not reproducible. It remains possible that survival in non-vegetative differentiated states could depend on the RRM, and this may explain why the RRM is more important in multicellular organisms (44,45). Alternatively, the contribution of the RRM to cellular processes may not be sufficient to detect a change in growth rate or cellular fitness, but could allow RRM-containing yeast to outcompete mutants defective in RNA-binding. This could have led to retention of the RRM over the course of evolution (8).

In contrast to yeast lacking the eIF4B RRM, we found that yeast lacking the NTD were highly sensitive to a number of conditions that WT cells are able to tolerate, and that for two of these conditions (urea and caffeine) effects conferred changes in translation in the mutant (Figure 2, S2, data not shown for caffeine). The mRNAs that showed decreased translation efficiency when the NTD was lacking had a number of features similar to those observed for an eIF4B null strain. The 5’UTRs of NTD-dependent mRNAs were longer and more structured than the average yeast mRNA (Figure 4), reinforcing many observations that eIF4B promotes translation of structured mRNAs (2,4,6,10). The mechanism by which eIF4B is proposed to promote translation of these mRNAs resides in its ability to interact with eIF4A and stimulate helicase activity. However, a report for direct interaction of these factors suggests that the 7-repeats domain of eIF4B binds eIF4A (18). In our study, we observed decreased translation of structured mRNAs when interaction of eIF4B with ribosomes was reduced by 80% upon NTD deletion (Figure 1, S1, Figure 4). Related components of the PIC, eIF4A and eIF4G, remained associated with ribosome fractions (Figure 1, S1). This suggests that the mechanism for eIF4B stimulation of structured mRNA translation resides at least to some extent in its ability to bind the ribosome (Figure 6). We have also previously reported defects in functional interaction of eIF4A and eIF4B when either the 7 repeats or the NTD is deleted, and observed that overexpression of Δntd has a dominant negative effect on an eIF4A mutant (13,17). Together these observations could indicate that deletion of the eIF4B NTD sequesters eIF4A in an inactive state off of the ribosome. However, we did not observe changes in the amount of eIF4A associated with small subunits and translating polysomes when eIF4B occupancy was decreased, arguing against this possibility and suggesting any direct interactions with eIF4A do not drive affinity for ribosome complexes. An alternative possibility is that deletion of the NTD prevents a PIC conformation required for optimal eIF4F activity. In either case, our data suggest the NTD of eIF4B mediates effective scanning through structured 5’UTRs while bound to the ribosome.

**Figure 6.**
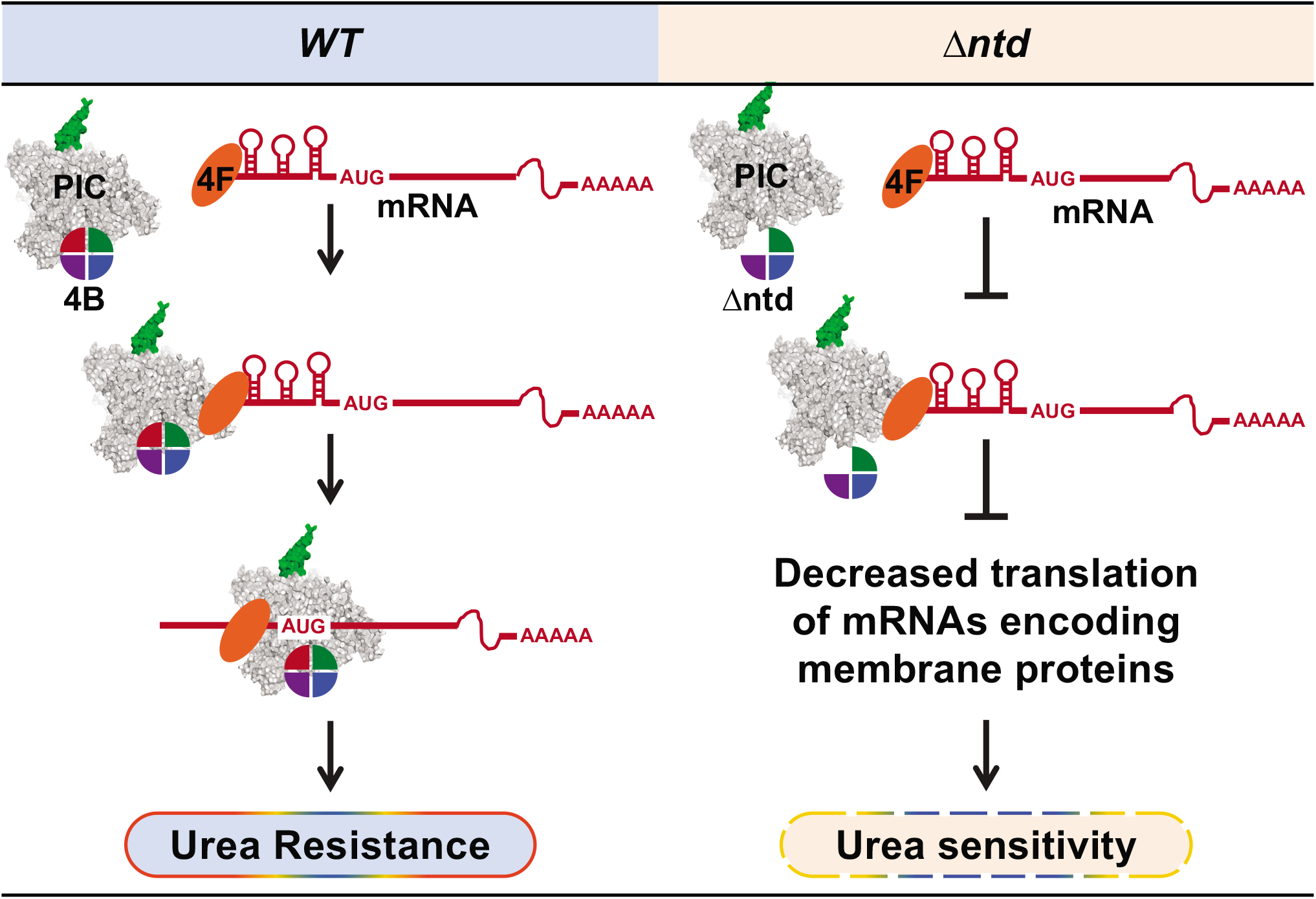
The NTD of eIF4B promotes translation of mRNAs with structured 5’-UTRs that reinforce the cellular periphery and allow a robust cellular response to urea. WT eIF4B promotes PIC loading and scanning of all mRNAs. Deletion of the NTD of eIF4B reduces translation of mRNAs with long structured 5’UTRs, indicating eIF4B promotes ribosome loading and scanning while bound to the PIC. Translation of these highly structured mRNAs are required to reconfigure the membrane proteome in response to stressors, providing urea resistance.

The strongest NTD-specific defect was observed in the presence of urea, which had no effect on WT growth rate or translation (P:M ratio) at concentrations that strongly repressed growth and translation of the mutant (Figure 2A-C). Urea affects several processes in *S. cerevisiae*, where it can serve as a nitrogen source, have effects on the membrane, and can denature structured nucleic acids. At the concentrations used here, it was reported that urea can readily cross the cellular membrane (37), presumably via the Dur3 transporter (46), and be used as a nitrogen source. Membrane blebbing and denaturation of nucleic acids is unlikely to occur at the ~0.5M urea used here for ribosome profiling (35,36). It is possible that gene expression changes are responsible for resistance at this level of urea, but upon deletion of eIF4B, the normal translation program is disrupted leading to sensitivity. Our RNASeq analysis indicates that in response to urea, there is a ~4-fold increase in DUR3 mRNA in WT cells, and a ~3-fold RNA increase in Δntd-expressing cells, suggesting a transcription-program or RNA stability difference (independent of eIF4B activity) is activated upon urea exposure, which leads to higher levels of *DUR3* mRNA. However, the *DUR3* mRNA is also dependent on the NTD for translation in urea, showing a 1.6-fold decreased TE in urea when the NTD was deleted relative to WT TE in urea (Table S6). It is possible that eIF4B-dependent translation of transmembrane urea transporters such as Dur3 can fine-tune the amount of urea imported or exported from the cells.

While most fungi use urease as a virulence factor and to break down urea, the Saccharomycetes (including *Saccharomyces, Candida*, and other genera) lack urease genes and instead use the Dur1,2 amidoylase to break down urea into ammonia and carbon dioxide in a biotin-dependent, two-step process (46,47). Our ribosome profiling data indicated that the *DUR1,2* mRNA was present at ~4-fold higher levels in both WT and mutant cells in response to urea (Table S6), but that the mRNA was translated ~1.5-fold less efficiently in urea when the NTD is deleted, indicating eIF4B-independent RNA control and eIF4B-dependent translational control of gene expression. Dur1,2 was demonstrated to be necessary for Candida differentiation and escape from macrophages during infection (48). There is very little sequence conservation between the NTD of yeast and human eIF4B, but these domains are well-conserved in *Saccharomyces* and *Candida* species. Because the *Saccharomyces* NTD deletion leads to changes in *DUR1,2* expression, the NTD of eIF4B could be a putative target for combating *Candida* infections.

Gene ontology analysis of ≥1.5-fold translation efficiency changes indicated enrichment in mRNAs encoding proteins associated with the membrane, and to a lesser extent cell wall, were translated at higher levels in WT cells in response to urea (Figure 3). Likewise, mRNAs encoding proteins associated with endomembrane system and modifications that arise within the ER and Golgi were unable to be translated efficiently in response to urea in the mutant cells (Figure 3, Table 1). The resulting membrane proteins are involved in a number of cellular processes. For instance, several proteins associated with adhesion during a-cell mating (Fig1, Fig2, Aga1, and Aga2; Table S6; Figure S4) showed TE changes in cells lacking the eIF4B NTD (decreases in TE in urea of 30-, 8-, 6- and 4-fold, respectively when compared to WT TE in urea). Recent analyses of uORF usage of yeast cells exposed to temperature shifts indicated that *AGA1* and *AGA2* showed changes in uORF usage in response to temperature shifts. In the presence of urea, or in response to NTD deletion we do not see changes in uORF usage of the *AGA2* mRNA. We do not see usage of the *AGA1* uORF in any of these experiments, suggesting the uORF occupancy observed in the former study was specific to changes in start codon fidelity in the high temperature response.

While we found that eIF4B promoted translation of mRNAs associated with specific gene ontology classes in response to urea, it may also be important that in response to stress, deletion of the NTD led to relative increases in translation of mRNAs associated with cytoplasmic translation and ribosome biogenesis, which lack structure in their 5’UTRs. In fact, we found substantial overlap in our analysis of gene ontology enrichment of unstructured mRNAs for eIF4B independence. In some conditions derepression of strong closed loop mRNAs that promote growth could be equally detrimental to cells as not producing membrane proteins needed for a particular stress response. We expect that this reprogramming is the result of ribosomes not being loaded onto eIF4B NTD-dependent mRNAs and therefore being more available for translation independent of eIF4B. It is also possible that these strong closed-loop mRNAs are recruited to ribosomes by a distinct pathway using eIF4G, which could explain why eIF4A and eIF4G occupancy with ribosomes were unaffected by NTD-deletion despite a reduction in cellular polysomes. These additional questions will be of great interest in future work.

## DATA AVAILABILITY

Raw sequencing reads and alignment files have been deposited in GEO with accession number GSE139097.

## ACKNOWLEDGEMENTS

The authors would like to thank Fujun Zhou for assistance generating yeast strains; Alan Hinnebusch and Jon Lorsch for providing reagents; Mary Thompson, Shardul Kulkarni, Audrey Michel, Onta Lin, Marie Saitou, Zhe Ji, and Michael Love for protocols and technical advice; and Paul Cullen, Joseph Barbi, and members of the Walker lab for helpful feedback on this manuscript.

## FUNDING

This work was supported by the National Institutes of Health [R00GM119173 to S.W.]; and start-up funds from the University at Buffalo College of Arts and Sciences.

## Notes

### Competing Interest Statement

The authors have declared no competing interest.

## REFERENCES

1. Mitchell, S.F., Walker, S.E., Rajagopal, V., Aitken, C.E. and Lorsch, J.R. (2011) Recruiting knotty partners: The roles of translation initiation factors in mRNA recruitment to the eukaryotic ribosome. Ribosomes: Structure, Function, and Dynamics, 155–169.

2. Ozes, A.R., Feoktistova, K., Avanzino, B.C. and Fraser, C.S. (2011) Duplex unwinding and ATPase activities of the DEAD-box helicase eIF4A are coupled by eIF4G and eIF4B. J Mol Biol, 412, 674–687.

3. Rozovsky, N., Butterworth, A.C. and Moore, M.J. (2008) Interactions between eIF4AI and its accessory factors eIF4B and eIF4H. RNA, 14, 2136–2148.

4. Dmitriev, S.E., Terenin, I.M., Dunaevsky, Y.E., Merrick, W.C. and Shatsky, I.N. (2003) Assembly of 48S translation initiation complexes from purified components with mRNAs that have some base pairing within their 5′ untranslated regions. Mol Cell Biol, 23, 8925–8933.

5. Mitchell, S.F., Walker, S.E., Algire, M.A., Park, E.H., Hinnebusch, A.G. and Lorsch, J.R. (2010) The 5′-7-methylguanosine cap on eukaryotic mRNAs serves both to stimulate canonical translation initiation and to block an alternative pathway. Mol Cell, 39, 950–962.

6. Sen, N.D., Zhou, F.J., Harris, M.S., Ingolia, N.T. and Hinnebusch, A.G. (2016) eIF4B stimulates translation of long mRNAs with structured 5 ′ UTRs and low closed-loop potential but weak dependence on eIF4G. P Natl Acad Sci USA, 113, 10464–10472.

7. Coppolecchia, R., Buser, P., Stotz, A. and Linder, P. (1993) A new yeast translation initiation factor suppresses a mutation in the eIF-4A RNA helicase. EMBO J, 12, 4005–4011.

8. Altmann, M., Muller, P.P., Wittmer, B., Ruchti, F., Lanker, S. and Trachsel, H. (1993) A Saccharomyces cerevisiae homologue of mammalian translation initiation factor 4B contributes to RNA helicase activity. EMBO J, 12, 3997–4003.

9. Andreou, A.Z. and Klostermeier, D. (2014) eIF4B and eIF4G jointly stimulate eIF4A ATPase and unwinding activities by modulation of the eIF4A conformational cycle. J Mol Biol, 426, 51–61.

10. Rogers, G.W., Jr., Richter, N.J., Lima, W.F. and Merrick, W.C. (2001) Modulation of the helicase activity of eIF4A by eIF4B, eIF4H, and eIF4F. J Biol Chem, 276, 30914–30922.

11. Sen, N.D., Zhou, F., Ingolia, N.T. and Hinnebusch, A.G. (2015) Genome-wide analysis of translational efficiency reveals distinct but overlapping functions of yeast DEAD-box RNA helicases Ded1 and eIF4A. Genome Res, 25, 1196–1205.

12. Park, E.H., Zhang, F., Warringer, J., Sunnerhagen, P. and Hinnebusch, A.G. (2011) Depletion of eIF4G from yeast cells narrows the range of translational efficiencies genome-wide. BMC Genomics, 12, 68.

13. Walker, S.E., Zhou, F., Mitchell, S.F., Larson, V.S., Valasek, L., Hinnebusch, A.G. and Lorsch, J.R. (2013) Yeast eIF4B binds to the head of the 40S ribosomal subunit and promotes mRNA recruitment through its N-terminal and internal repeat domains. RNA, 19, 191–207.

14. Fleming, K., Ghuman, J., Yuan, X., Simpson, P., Szendroi, A., Matthews, S. and Curry, S. (2003) Solution structure and RNA interactions of the RNA recognition motif from eukaryotic translation initiation factor 4B. Biochemistry, 42, 8966–8975.

15. Xue, B., Dunbrack, R.L., Williams, R.W., Dunker, A.K. and Uversky, V.N. (2010) PONDR-FIT: a meta-predictor of intrinsically disordered amino acids. Biochim Biophys Acta, 1804, 996–1010.

16. Niederberger, N., Trachsel, H. and Altmann, M. (1998) The RNA recognition motif of yeast translation initiation factor Tif3/eIF4B is required but not sufficient for RNA strand-exchange and translational activity. RNA, 4, 1259–1267.

17. Zhou, F.J., Walker, S.E., Mitchell, S.F., Lorsch, J.R. and Hinnebusch, A.G. (2014) Identification and characterization of functionally critical, conserved motifs in the internal repeats and N-terminal domain of yeast translation initiation factor 4B (yeIF4B). Journal of Biological Chemistry, 289, 11860–11860.

18. Andreou, A.Z., Harms, U. and Klostermeier, D. (2017) eIF4B stimulates eIF4A ATPase and unwinding activities by direct interaction through its 7-repeats region. RNA Biol, 14, 113–123.

19. Park, E.H., Walker, S.E., Zhou, F., Lee, J.M., Rajagopal, V., Lorsch, J.R. and Hinnebusch, A.G. (2013) Yeast eukaryotic initiation factor 4B (eIF4B) enhances complex assembly between eIF4A and eIF4G in vivo. J Biol Chem, 288, 2340–2354.

20. Altmann, M., Wittmer, B., Methot, N., Sonenberg, N. and Trachsel, H. (1995) The Saccharomyces cerevisiae translation initiation factor Tif3 and its mammalian homologue, eIF-4B, have RNA annealing activity. EMBO J, 14, 3820–3827.

21. Mulleder, M., Capuano, F., Pir, P., Christen, S., Sauer, U., Oliver, S.G. and Ralser, M. (2012) A prototrophic deletion mutant collection for yeast metabolomics and systems biology. Nat Biotechnol, 30, 1176–1178.

22. Lee, M.E., DeLoache, W.C., Cervantes, B. and Dueber, J.E. (2015) A Highly Characterized Yeast Toolkit for Modular, Multipart Assembly. ACS Synth Biol, 4, 975–986.

23. Bochner, B.R., Gadzinski, P. and Panomitros, E. (2001) Phenotype microarrays for high-throughput phenotypic testing and assay of gene function. Genome Res, 11, 1246–1255.

24. Lee, B., Udagawa, T., Singh, C.R. and Asano, K. (2007) Yeast phenotypic assays on translational control. Methods Enzymol, 429, 105–137.

25. Herrmannova, A., Prilepskaja, T., Wagner, S., Sikrova, D., Zeman, J., Poncova, K. and Valasek, L.S. (2020) Adapted formaldehyde gradient cross-linking protocol implicates human eIF3d and eIF3c, k and l subunits in the 43S and 48S pre-initiation complex assembly, respectively. Nucleic Acids Res, 48, 1969–1984.

26. Liu, X., Schuessler, P.J., Sahoo, A. and Walker, S.E. (2019) Reconstitution and analyses of RNA interactions with eukaryotic translation initiation factors and ribosomal preinitiation complexes. Methods, 162-163, 42–53.

27. Ingolia, N.T., Ghaemmaghami, S., Newman, J.R. and Weissman, J.S. (2009) Genome-wide analysis in vivo of translation with nucleotide resolution using ribosome profiling. Science, 324, 218–223.

28. McGlincy, N.J. and Ingolia, N.T. (2017) Transcriptome-wide measurement of translation by ribosome profiling. Methods, 126, 112–129.

29. Guydosh, N.R. and Green, R. (2014) Dom34 rescues ribosomes in 3′ untranslated regions. Cell, 156, 950–962.

30. Michel, A.M., Mullan, J.P., Velayudhan, V., O’Connor, P.B., Donohue, C.A. and Baranov, P.V. (2016) RiboGalaxy: A browser based platform for the alignment, analysis and visualization of ribosome profiling data. RNA Biol, 13, 316–319.

31. Martin, M. (2011) Cutadapt Removes Adapter Sequences From High-Throughput Sequencing Reads. EMBnet.journal, 17, 10–12.

32. Langmead, B., Trapnell, C., Pop, M. and Salzberg, S.L. (2009) Ultrafast and memory-efficient alignment of short DNA sequences to the human genome. Genome Biol, 10, R25.

33. Love, M.I., Huber, W. and Anders, S. (2014) Moderated estimation of fold change and dispersion for RNA-seq data with DESeq2. Genome Biol, 15, 550.

34. Thompson, M.K., Rojas-Duran, M.F., Gangaramani, P. and Gilbert, W.V. (2016) The ribosomal protein Asc1/RACK1 is required for efficient translation of short mRNAs. Elife, 5.

35. Lambert, D. and Draper, D.E. (2012) Denaturation of RNA secondary and tertiary structure by urea: simple unfolded state models and free energy parameters account for measured m-values. Biochemistry, 51, 9014–9026.

36. Necas, O. and Svoboda, A. (1973) Effect of urea on the plasma membrane particles in yeast cells and protoplasts. Protoplasma, 77, 453–466.

37. Cooper, T.G. and Sumrada, R. (1975) Urea transport in Saccharomyces cerevisiae. J Bacteriol, 121, 571–576.

38. Dolan, K., Montgomery, S., Buchheit, B., Didone, L., Wellington, M. and Krysan, D.J. (2009) Antifungal activity of tamoxifen: in vitro and in vivo activities and mechanistic characterization. Antimicrob Agents Chemother, 53, 3337–3346.

39. Kim, J., Oh, J. and Sung, G.H. (2016) Regulation of MAP kinase Hog1 by calmodulin during hyperosmotic stress. Biochim Biophys Acta, 1863, 2551–2559.

40. Kertesz, M., Wan, Y., Mazor, E., Rinn, J.L., Nutter, R.C., Chang, H.Y. and Segal, E. (2010) Genome-wide measurement of RNA secondary structure in yeast. Nature, 467, 103–107.

41. Costello, J., Castelli, L.M., Rowe, W., Kershaw, C.J., Talavera, D., Mohammad-Qureshi, S.S., Sims, P.F.G., Grant, C.M., Pavitt, G.D., Hubbard, S.J. et al. (2015) Global mRNA selection mechanisms for translation initiation. Genome Biol, 16.

42. Methot, N., Song, M.S. and Sonenberg, N. (1996) A region rich in aspartic acid, arginine, tyrosine, and glycine (DRYG) mediates eukaryotic initiation factor 4B (eIF4B) self-association and interaction with eIF3. Mol Cell Biol, 16, 5328–5334.

43. Methot, N., Pickett, G., Keene, J.D. and Sonenberg, N. (1996) In vitro RNA selection identifies RNA ligands that specifically bind to eukaryotic translation initiation factor 4B: the role of the RNA remotif. RNA, 2, 38–50.

44. Naranda, T., Strong, W.B., Menaya, J., Fabbri, B.J. and Hershey, J.W. (1994) Two structural domains of initiation factor eIF-4B are involved in binding to RNA. J Biol Chem, 269, 14465–14472.

45. Methot, N., Pause, A., Hershey, J.W. and Sonenberg, N. (1994) The translation initiation factor eIF-4B contains an RNA-binding region that is distinct and independent from its ribonucleoprotein consensus sequence. Mol Cell Biol, 14, 2307–2316.

46. Navarathna, D.H., Das, A., Morschhauser, J., Nickerson, K.W. and Roberts, D.D. (2011) Dur3 is the major urea transporter in Candida albicans and is co-regulated with the urea amidolyase Dur1,2. Microbiology, 157, 270–279.

47. Navarathna, D.H., Harris, S.D., Roberts, D.D. and Nickerson, K.W. (2010) Evolutionary aspects of urea utilization by fungi. FEMS Yeast Res, 10, 209–213.

48. Ghosh, S., Navarathna, D.H., Roberts, D.D., Cooper, J.T., Atkin, A.L., Petro, T.M. and Nickerson, K.W. (2009) Arginine-induced germ tube formation in Candida albicans is essential for escape from murine macrophage line RAW 264.7. Infect Immun, 77, 1596–1605.

