## Supplementary Materials for "The N-terminal domain of Eukaryotic Initiation Factor 4B Drives Yeast Translational Control in Response to Urea"

**Supplementary Table 1: Yeast strains used in this study**

| Strain Name | Genotype |
|---|---|
| YSW4 | *Mata, his3Δ1, leu2Δ0, ura3Δ0, met15Δ0, tif3Δ0* |
| YSW5 | *Mata, his3Δ1, leu2Δ0, ura3Δ0, met15Δ0, tif3Δ0 [pSW150:HIS3, LEU2, URA3, MET15, TIF3]* |
| YSW6 | *Mata, his3Δ1, leu2Δ0, ura3Δ0, met15Δ0, tif3Δ0 [pSW151:HIS3, LEU2, URA3, MET15, tif3Δntd]* |
| YSW7 | *Mata, his3Δ1, leu2Δ0, ura3Δ0, met15Δ0, tif3Δ0 [pSW152:HIS3, LEU2, URA3, MET15, tif3Δrrm]* |
| YSW273 | *Mata, his3Δ1, leu2Δ0, ura3Δ0, met15Δ0, tif3Δ0 [pSW150:HIS3, LEU2, URA3, MET15, TIF3] [pAS48:KanR, fig2 UTR+1st 30-GFP]* |
| YSW274 | *Mata, his3Δ1, leu2Δ0, ura3Δ0, met15Δ0, tif3Δ0 [pSW150:HIS3, LEU2, URA3, MET15, TIF3] [pAS49:KanR, vba2 UTR+1st 30-GFP]* |
| YSW277 | *Mata, his3Δ1, leu2Δ0, ura3Δ0, met15Δ0, tif3Δ0 [pSW151:HIS3, LEU2, URA3, MET15, tif3Δntd] [pAS48:KanR, fig2 UTR+1st 30-GFP]* |
| YSW278 | *Mata, his3Δ1, leu2Δ0, ura3Δ0, met15Δ0, tif3Δ0 [pSW151:HIS3, LEU2, URA3, MET15, tif3Δntd] [pAS49:KanR, vba2 UTR+1st 30-GFP]* |

**Supplementary Table 2: Plasmids used in this study**

| Addgene ID | Plasmid Name | Relevant Gene(s) | Parental Vector | Markers | Template/parts vectors | Primers used |
|---|---|---|---|---|---|---|
| 168033 | pSW150 | *TIF3-His6* | pHLUM | His, Leu, Ura, Met | pFJZ056 | SW223/224 |
| 168034 | pSW151 | *tif3Δntd-His6* | pHLUM | His, Leu, Ura, Met | pFJZ141 | SW223/224 |
| 168035 | pSW152 | *tif3Δrrm-His6* | pHLUM | His, Leu, Ura, Met | pFJZ140 | SW223/224 |
| N/A | pAS39 | *FIG2-P-5'UTR-First 30-Venus* | pYTK001 | E. coli CamR | BY4741 DNA | AS160/171 |
| N/A | pAS40 | *VBA2-P-5'UTR-First 30-Venus* | pYTK001 | E. coli CamR | BY4741 DNA | AS164/170 |
| N/A | pAS45 | *Intermediate vector for golden gate cloning* | pYTK003 | Yeast KanR<br>E. coli AmpR | pYTK047<br>pYTK068<br>pYTK077<br>pYTK081<br>pYTK083 | N/A |
| 168036 | pAS48 | *FIG2-P-5'UTR-First 30-Venus* | pAS45 | Yeast KanR<br>E. coli AmpR | pAS39<br>pYTK045<br>pYTK053 | N/A |
| 168037 | pAS49 | *VBA2-P-5'UTR-First 30-Venus* | pAS45 | Yeast KanR<br>E. coli AmpR | pAS40<br>pYTK045<br>pYTK053 | N/A |

**Supplementary Table 3: Primers used in this study**

| Primer (cloning) | Sequence |
|---|---|
| SW223 | GGCCGCTCTAGAACTAGTGACCTAATTGACACCGTAC |
| SW224 | CGACGTAGTCGAGGATCGCTTCTTCTTTTGAATATTACC |
| AS160 | TTTCGTCTCGTCGGGGTCTCGAACGCGAGCCTTCCCTTTCAGTA |
| AS171 | TTTCGTCTCGGGTCGGTCTCGAGAACCACTATTCTGGTAGAACACAATTT |
| AS164 | TTTCGTCTCGTCGGGGTCTCGAACGTGGCAACCACATTCTAAG |
| AS170 | TTTCGTCTCGGGTCGGTCTCGAGAACCACCAAGTGCATCAGAAAATG |

| Primer (RT) | Sequence |
|---|---|
| ACT1/YFL039C | CGTCTGGATTGGTGGTTCTATC, GGACCACTTTCGTCGTATTCTT |
| ADH4/YGL256W | TGCTGTCAACGATCCATCTAC, AGAGGCGGTGGAAACATAAG |
| AGA1/YNR044W | GTAAGTGAAGCCACGAGTACAT CAGACAAGGAGGAGGATGAAAG |
| AGA2/YGL032C | GAATCGACGCCGTACTCTTT, GGGTGAGAACCGCAATTACT |
| FIG2/YCR089W | GCAGTGTTGACAGGTTTGTTT, CTTATGGTGGTGGTAGCAGTAG |
| FLUC | TCATCATGGACAGCAAGACC, CACGAAGTCGTACTCGTTGAA |
| GCN4/YEL009C | CCAGTTACCACTGACGATGTT, AGTTGTCGAGACTTCCAGATTG |
| VBA2/YBR293W | CCGAACTGGTTGATTGGTCTAT, GCAGTAGCTTGGTCACTCTTAG |
| YER186C | GTTTACCAGAGGTGGGTGATAG, CCTCCTTAGTTCTGCACATACA |
| YOR015W | GCATACTTCAATCGATGTGTCTTC, AAATAGAGGAAAGGCAGAAGGAA |

**Supplementary Table 4:**
**Barcode information for RiboSeq and RNASeq library preparation**

**(attached in a separate file)**

**Supplementary Table 5:**

**Stress conditions that significantly affect the growth of the NTD deletion strain**

| Chemical | Panel | Average Height Difference (Δ*ntd* vs WT) |
|---|---|---|
| 3% Urea | Osmolytes | -129 |
| Trp-Ser | Peptide Nitrogen sources | -113 |
| Trp-Asp | Peptide Nitrogen sources | -112 |
| 4% Urea | Osmolytes | -109 |
| Trp-Gly | Peptide Nitrogen sources | -105 |
| Trp-Ala | Peptide Nitrogen sources | -103 |
| Tyr-Ala | Peptide Nitrogen sources | -101 |
| Pro-Gln | Peptide Nitrogen sources | -100 |
| Tyr-Leu | Peptide Nitrogen sources | -97 |
| Lys-Ala | Peptide Nitrogen sources | -97 |
| Thiourea | Phosphorus and Sulfur Sources | -94 |
| 2% Urea | Osmolytes | -93 |
| Apramycin sulfate | Chemical Sensitivity | -93 |
| Tamoxifen | Chemical Sensitivity | -93 |
| Phe-Ala | Peptide Nitrogen sources | -92 |
| Chromium (III) chloride | Chemical Sensitivity | -89 |
| L-Citrulline | Nitrogen Sources | -89 |
| Tyr-Gln | Peptide Nitrogen sources | -89 |
| 6% Potassium chloride | Osmolytes | -88 |
| Met-Arg | Peptide Nitrogen sources | -87 |
| Ser-Met | Peptide Nitrogen sources | -85 |
| Cystathionine | Phosphorus and Sulfur Sources | -83 |
| Trp-Leu | Peptide Nitrogen sources | -83 |
| L-Cysteine | Phosphorus and Sulfur Sources | -82 |
| Met-Leu | Peptide Nitrogen sources | -82 |
| 5% Potassium Chloride | Osmolytes | -82 |
| Cobalt (II) chloride | Chemical Sensitivity | -82 |
| m-Inositol | Nutrient Supplements | -82 |
| Met-Gln | Peptide Nitrogen sources | -81 |
| Poly-L-lysine | Chemical Sensitivity | -81 |
| Thr-Arg | Peptide Nitrogen sources | -80 |
| Caffeine | Chemical Sensitivity | -80 |
| Cysteamine | Phosphorus and Sulfur Sources | -80 |

**Supplementary Table 6:**

**Comparison of RNAseq, Riboseq and Translation efficiency**

**(attached in a separate file)**

**Supplementary Table 7: Gene ontology enrichment for mRNAs with ≥1.5-fold increased translational efficiency in ∆*ntd* in response to urea**

| GO term | p-Value |
|---|---|
| Cytoplasm | 5.39E-07 |
| Oxidation-reduction process | 4.13E-05 |
| Energy reserve metabolic process associated | 1.54E-03 |
| Mitochondrion | 5.83E-03 |
| Protein refolding | 1.24E-02 |
| Mitochondrial envelope | 1.94E-02 |
| Response to temperature stimulus associated | 2.05E-02 |
| Cellular response to chemical stimulus | 2.32E-02 |
| Cellular response to oxidative stress | 2.62E-02 |
| Carbohydrate metabolic process associated | 3.18E-02 |
| Purine ribonucleoside triphosphate metabolic process | 4.45E-02 |
| Oxidoreductase activity associated | 4.89E-02 |

**Supplementary Table 8: Gene ontology enrichment for mRNAs with ≥1.5-fold increased translational efficiency in ∆*ntd* without urea**

| GO term | p-Value |
|---|---|
| Cytosolic ribosome | 1.26E-35 |
| Cytoplasmic translation | 2.66E-30 |
| Ribonucleoprotein complex | 8.17E-11 |
| Translational elongation | 4.44E-10 |
| Organonitrogen compound biosynthetic process | 2.99E-08 |
| Cytosol | 5.81E-08 |
| Cellular amide biosynthetic process | 2.57E-07 |
| Translation | 4.16E-07 |
| Peptide biosynthetic process | 5.44E-07 |
| Organonitrogen compound metabolic process | 2.08E-06 |
| Peptide metabolic process | 3.99E-06 |
| Translational frameshifting | 2.76E-03 |
| Non-membrane-bounded organelle | 3.63E-03 |
| Carboxylic acid metabolic process | 7.57E-03 |

**Supplementary Table 9: Gene ontology enrichment for mRNAs with ≥1.5-fold decreased translational efficiency in ∆*ntd* without urea**

| GO term | p-Value |
|---|---|
| Intracellular membrane-bounded organelle | 5.95E-03 |
| Membrane-bounded organelle | 8.32E-03 |

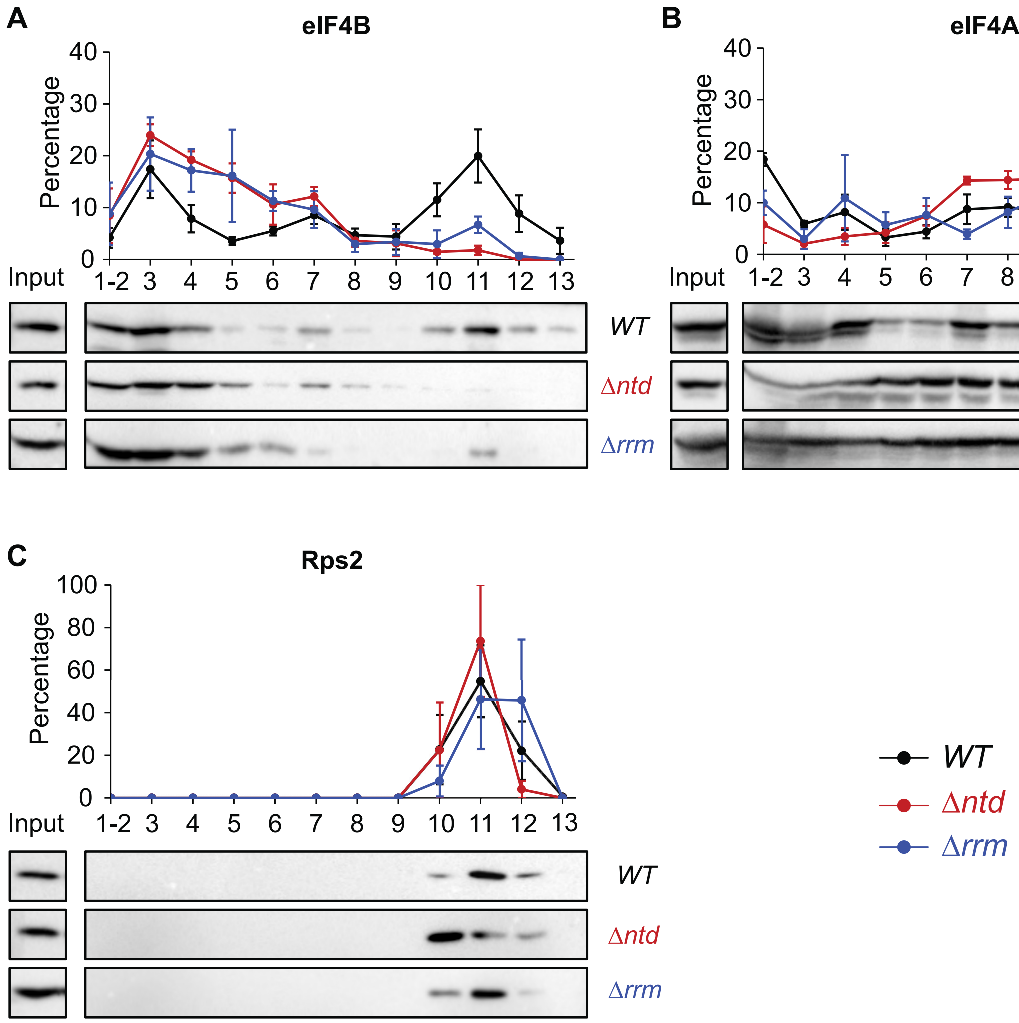


**Supplementary Figure 1: eIF4B comigrates with both small subunits and mRNPs in gradients following formaldehyde crosslinking.**

Strains harboring WT (black), Δntd (red), and Δrrm (blue) eIF4B were grown in SD media prior to formaldehyde crosslinking. Protein precipitates from lysates were separated on 7.5-30% sucrose gradients and blotted for eIF4B-His6 (A), yeast eIF4A (B), or small subunit protein Rps2 (C).

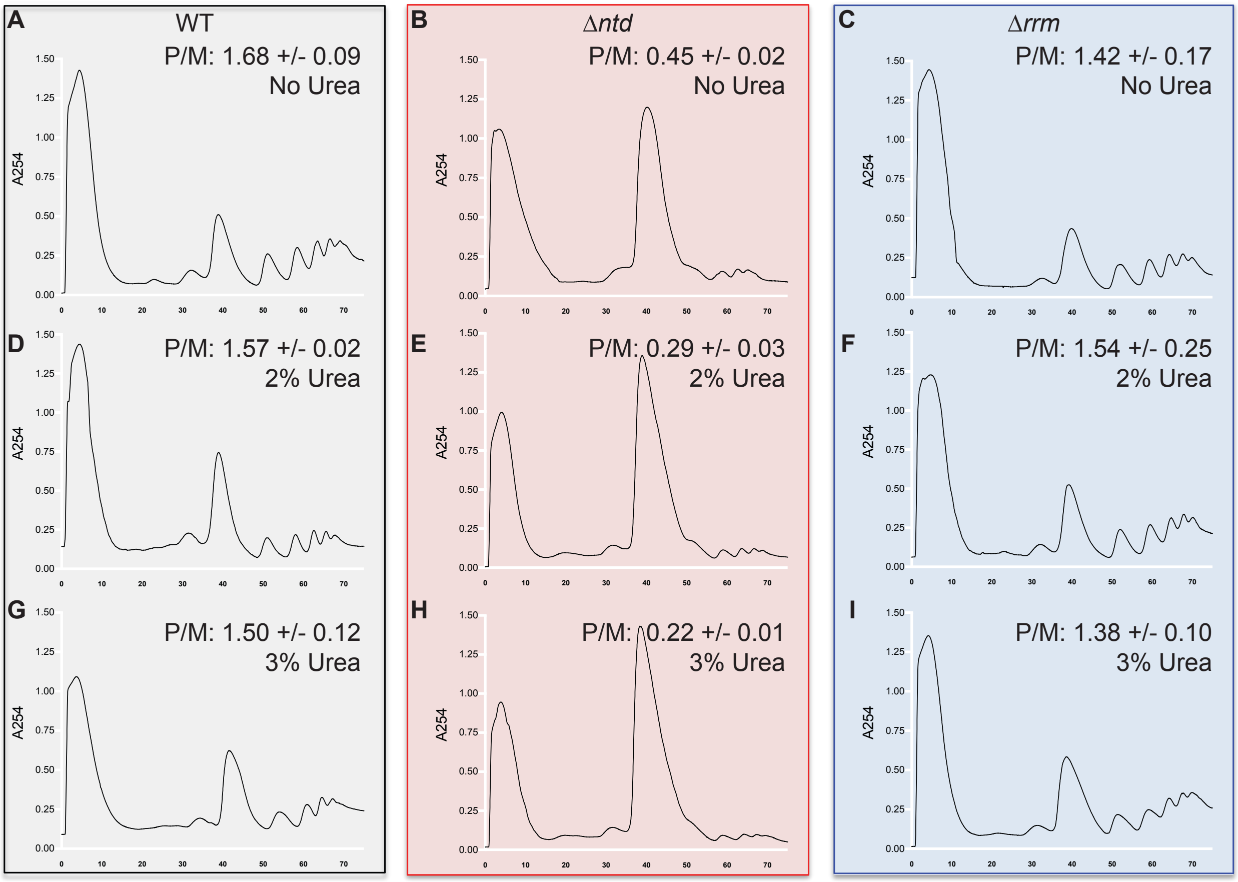


**Supplementary Figure 2: Polysome to monosome ratios show both the NTD of eIF4B and Urea conferred changes in the number of ribosomes loaded per mRNA.**

Representative sucrose gradient analysis for calculating polysome to monosome changes for no urea (A-C) 2% urea (D-F) and 3% Urea (G-I) in cells harboring WT eIF4B (A, D, G), Δntd (B, E, H), or Δrrm (C, F, I). Deletion of the NTD globally repressed translation, while addition of urea conferred further decreases. The low level of translating ribosomes in the Δntd mutant makes the increase in monosomes upon urea addition more apparent by eye than the reduction in polysomes.

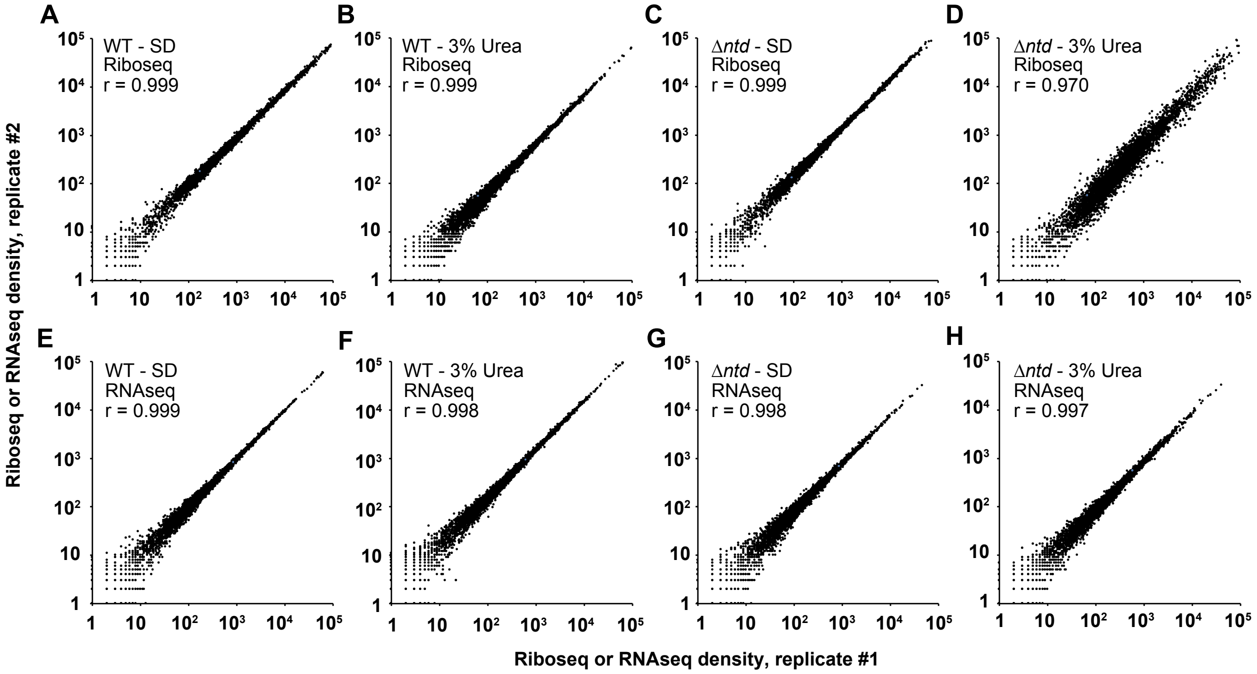


**Supplementary Figure 3: Riboseq and RNAseq reads are reproducible for two independent cultures of WT and NTD deletion mutant with and without urea.**

Riboseq (A-D) and RNAseq (E-H) densities on individual mRNAs were compared for two biological replicates of WT and NTD deletion mutants with and without urea to determine reproducibility of samples. Pearson correlation coefficients (r) were calculated from each comparison plot.

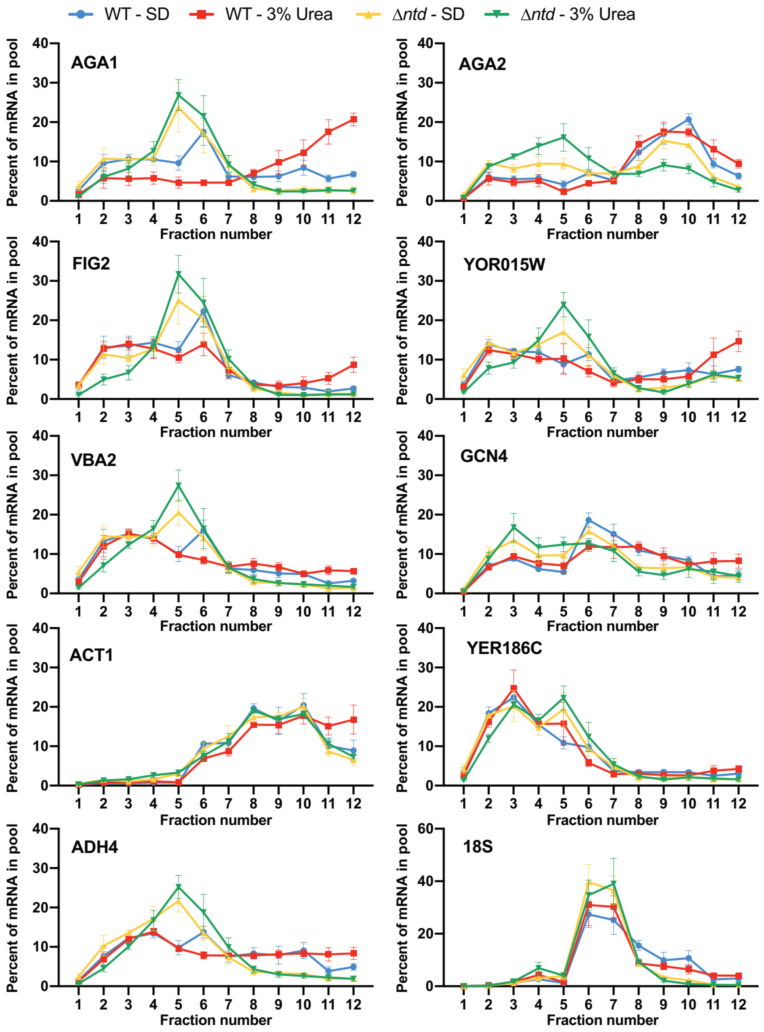


**Supplementary Figure 4: The NTD of eIF4B and exposure to Urea promote mRNA-specific changes in translation.** Polysome qRT-PCR showing association of representative mRNAs with polysomes (Fraction No. 8 to 12) in four different conditions: WT without urea (blue), WT with 3% urea (red), NTD deletion mutant (Δ*ntd*) without urea (yellow), and Δ*ntd* with 3% urea (green). Values were normalized to the RNA spike-in control in each fraction and plotted as the percentage of RNA in all fractions. Results from three biological replicates ± SEM are shown. The mRNAs showing increased association with polysome-containing fractions (fractions 8-12) in WT in response to 3% urea (red) are: *AGA1*, *FIG2*, and *YOR015W*; while genes showing decreased association with polysome-containing fractions as a result of NTD-deletion (green and yellow) are *AGA1*, *FIG2*, *YOR015W*, *VBA2*, and *ADH4*. The 18S also shows decreased levels in polysomes as a function of urea and the NTD, and verifies the location of subunits and ribosomes. *AGA2* shows decreased polysome association upon NTD deletion and further decreases following urea exposure.

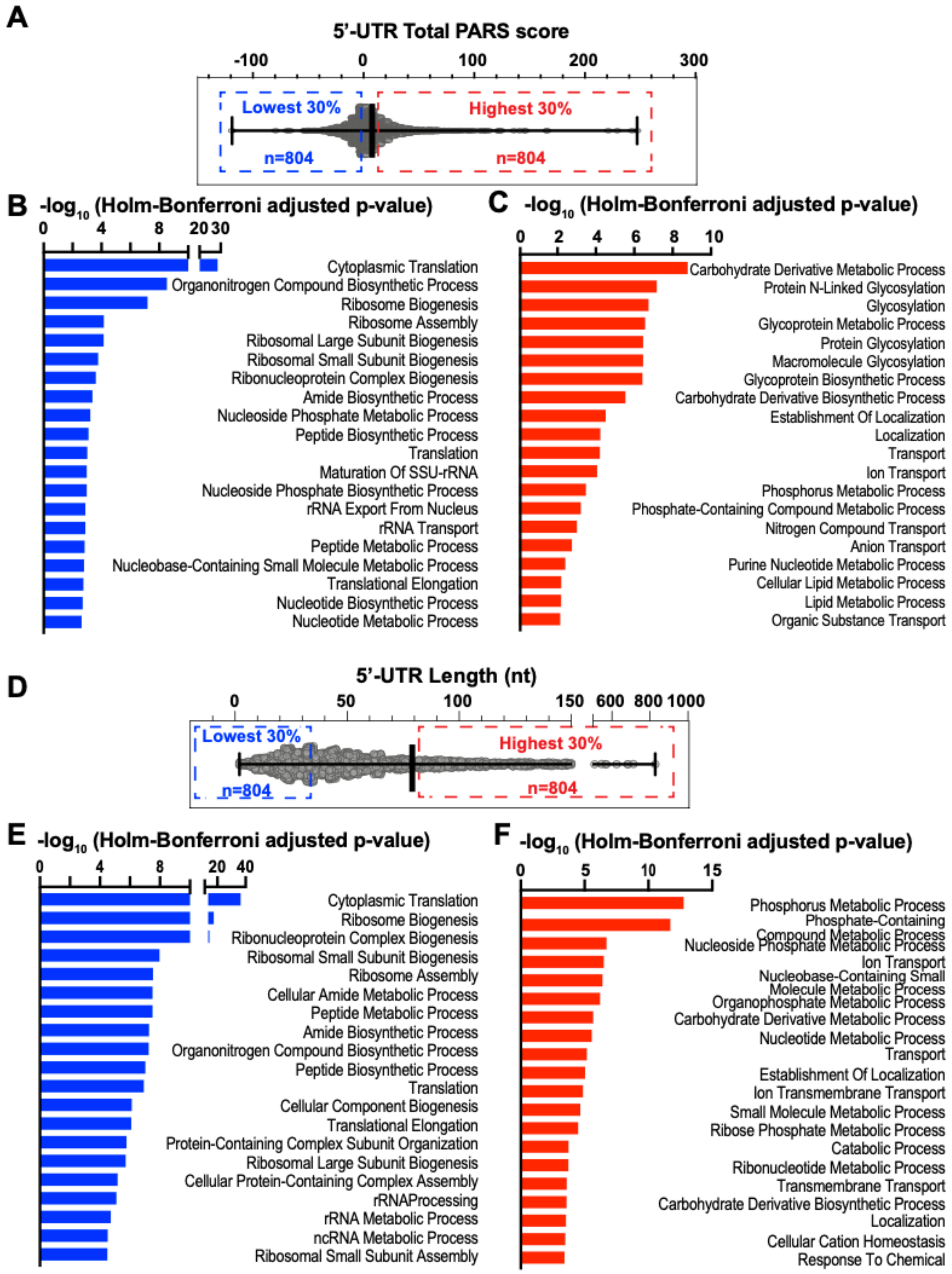


**Supplementary Figure 5: Gene ontology analysis for yeast mRNAs grouped based on 5'UTR length (D-F) and propensity for involvement in secondary structure (A-C.)** Total 5'UTR PARS scores (A) and Lengths of 5'UTRs (D) for all yeast mRNAs were ranked, and the 30% highest (C, F) and lowest (B, E) for each ranking were analyzed for gene ontology enrichment at Yeastmine.
